# Chemical activity profiling reveals how exposure to drugs or dietary compounds alters gut microbial biotransformation capacity

**DOI:** 10.1101/2025.05.20.655107

**Authors:** Hélène Bigonne, Shuhuan Zhai, Jacob Folz, Katherine Hurley, Aishwarya Murali, Franziska Maria Zickgraf, Hennicke Kamp, Bennard van Ravenzwaay, Georg Aichinger, Shana J. Sturla

**Author notes:** Equal contributions (Hélène Bigonne, Shuhuan Zhai).

## Abstract

The gut microbiome catalyzes biotransformation reactions that influence intestinal absorption as a basis of microbiome-host interactions. A better understanding of microbiota biotransformation capacity, and its alteration in dysregulated states, would enable the prediction of individual responses to drugs and toxins and improve safety assessment. Here, we profiled chemical activities in rat gut microbiota ex vivo, and quantified biotransformation capacity changes induced by oral exposure to eight drugs and dietary compounds: tobramycin, colistin, acesulfame potassium, saccharin, bovine serum albumin (BSA), meropenem, doripenem, and daidzein. We implemented an approach involving inoculation with metabolic probes during microbiota fermentations and measured their degradation. In most exposure groups, we observed no alteration of microbiota biotransformation capacity, however, we detected significant alterations in biotransformation rates after exposure to meropenem, doripenem and tobramycin. Interestingly, common patterns of biotransformation capacity were observed in the gut microbiomes from rats exposed to carbapenems and partially shared in microbiomes exposed to tobramycin. These results aligned well with prior metagenomic and metabolomic findings. Further, correlations between microbial taxa and reaction rates were assessed to establish a link between specific bacteria and probe degradation. This functionally relevant strategy revealed alterations of microbiota biotransformation capacity, as induced by in vivo exposures.

## Introduction

Mammals and their gut microbiota have co-evolved to form an intricate and mutually beneficial relationship. Reflecting distinct evolutionary pressures, the diverse microbial communities that constitute the gut microbiota carry out a wide array of chemical reactions, often reversing host metabolism (Koppel et al., 2017). Enzymes expressed in this complex ecosystem alter chemical structures of both endogenous substrates and xenobiotic compounds such as drugs and dietary compounds, ultimately affecting their biological function or toxicity (Collins & Patterson, 2020; Koppel et al., 2017). This biotransformation capacity, also referred to as gut microbial functionality, has a profound influence on the metabolism, nutrition, physiology, immune function, and mental well-being of its host (Appleton, 2018; Bull & Plummer, 2014; Fan & Pedersen, 2021). Consequently, complex interactions over the lifespan, including lifestyle, age, diet, disease, and medication can disrupt the host-microbiota interplay (Clarke et al., 2019; Nicholson et al., 2012), and lead to dysbiosis (an imbalance in gut microbial composition and biotransformation capacity) which in turn contributes to disorders such as irritable bowel syndrome, non-alcoholic liver disease, obesity, type 2 diabetes, and malnutrition (Carding et al., 2015; Fan & Pedersen, 2021; Rooks & Garrett, 2016). Beyond alterations in bacterial composition, understanding microbiota dysbiotic states by assessing its biotransformation capacity is necessary to establish biomarkers of health and disease and incorporate the gut microbiome into xenobiotic risk assessment (Fan & Pedersen, 2021; Isenring et al., 2023; Lacroix et al., 2015; Noecker & Turnbaugh, 2024; Stevanoska et al., 2024). Indeed, analysis of biotransformation capacities is crucial to capture the biochemical consequences of microbiota community disruptions, as it directly reflects how these alterations impact metabolic pathways at the ecosystem-level.

Assessing gut microbiome biotransformation capacity, including its alterations in dysbiotic states, presents a significant challenge. To tackle this, common approaches combine metabolomics and metagenomics. For instance, strong associations of gut microbial composition, fecal metabolome and visceral fat mass were revealed in a study based on the analysis of 1,116 fecal metabolites from a population of 786 female twins (from the TwinsUK cohort) (Zierer et al., 2018). Additionally, metabolism of specific substances could be predicted from metagenomic and genomic sequence data in gnotobiotic mice (Zimmermann et al., 2019). Furthermore, this combination of metabolomic and metagenomic analyses is increasingly employed to characterize alterations in the gut microbiome, particularly following exposure to drugs or dietary chemicals (Murali et al., 2021; Murali et al., 2022; Murali, Giri, et al., 2023; Murali, Zickgraf, et al., 2023). However, accurately predicting the biotransformation capacity based on microbiota metagenomic data is not currently entirely possible in humans. This was highlighted by a recent analysis of 1,135 individuals from a Dutch population-based cohort, in which no correlation was found between the levels of microbiome-derived metabolites in plasma and feces and the abundance of the corresponding metabolic genes (Pascal Andreu et al., 2023). While metabolomics provides a snapshot of the metabolic consequences of gut microbiome alterations, it does not directly assess the microbiome’s functional capacity to carry out specific biotransformation reactions, or how these are altered by perturbations.^

Chemical activity profiling offers a direct assessment of a microbiome’s biotransformation capacity. van de Steeg et al. (2018) introduced the concept of using a set of diverse compounds specifically selected for their known degradation by the gut microbiota, such that when incubated together with pooled microbiota samples from healthy donors, they acted as probes, and their degree of degradation after 24 h indicated the occurrence of particular biotransformation reactions. The expected reactions were further validated by the targeted detection of corresponding metabolites, ultimately allowing for the simultaneous assessment of the microbiome’s functional capacities. In a recent milestone study, Zimmerman et al. evaluated 76 bacterial taxa (representing the major phyla of the human gut microbiome) for their capacity to chemically modify 271 drugs in 12-h incubations of each 20,596 drug-bacteria pairs (Zimmermann et al., 2019). While both approaches allow the observation of biotransformation reactions carried out by the microbiota, quantitative metrics are still lacking for the comparison of changes in biotransformation capacities between different altered states of the microbiome. This last point was addressed by Culp et al., in a study mapping metabolic activities in 29 healthy human gut microbiomes, each individually incubated for 48 h with one of 22 dietary xenobiotics (Culp et al., 2024). LC-MS analysis of the resulting metabolites revealed a wide range of biotransformation reactions, including ester bond cleavage, deglycosylation, ring-opening, double bond reduction, demethylation, dehydroxylation, and methoxy group cleavage. However, such alterations in biotransformation capacities have not yet been characterized by chemical activity profiling in microbiota altered in vivo, due to chemical or drug exposures.

In the present study, we characterized biotransformation capacity of rat gut microbiomes and compared their alteration following exposure to various xenobiotics. After 4 weeks of exposure, we collected fecal microbiota samples and quantified the degradation rates of probes in ex vivo fermentations. Twelve probes representing distinct microbiota-mediated reactions were used to profile the biotransformation capacity of the rat gut microbiomes. Fecal metabolomics and metagenomic data from the same animals were also collected, providing an integrated, multi-omic assessment of xenobiotic-induced microbiome alterations.

## Materials and methods

### General considerations

Chemical and reagent sources are listed in **Table S1**. For the preparation of Macfarlane media, ultrapure water supplied from a Milli-Q IQ 7000 purification system (Merck) was used. In all other cases, LCMS-grade water was used.

### Animal experiments

Two 28-day oral toxicity studies were carried out on Wistar rats [CrI/WI(Han)] of 70±1 days old, as previously described in Murali et al. (2022); Murali, Giri, et al. (2023); Murali, Zickgraf, et al. (2023) (Sampling Groups 1 and 2, Figure 1). Both were approved by the local authorizing agency for animal experiments (Landesuntersuchungsamt Rheinland-Pfalz, Koblenz, Germany, approval number 23,177–07/G 18-3-098) and performed under the OECD 407 Principles of Good Laboratory Practice guidelines and the GLP provisions of the German Chemicals Act. In a first sampling, rats were administered antibiotics (the aminoglycoside tobramycin and the polymyxin E colistin sulfate (colistin)) (Murali, Giri, et al., 2023), low-calorie artificial sweeteners (acesulfame potassium (acesulfame K) and saccharin) (Murali et al., 2022) or bovine serum albumin (BSA), a commonly used vehicle. In the second sampling, rats were administered the broad-spectrum carbapenem antibiotics meropenem trihydrate (meropenem) and doripenem hydrate (doripenem) (Murali, Zickgraf, et al., 2023) and the isoflavone daidzein. Two dose levels were used, defined as low dose (LD) and high dose (HD); these were set based on available literature data balancing evidence for altering the targeted microbiome population without causing systemic toxicity (**Table S2**). Equal numbers of male and female rats were used and there were corresponding control animals per exposure and per sex. Stratification and population of each group are summarized in **Table S3**. The sampling timeline is presented in **Figure 1**. Samples collected on day 28 were used for DNA isolation, 16S gene PCR amplification and sequencing, as described in Murali et al. (2022); Murali, Giri, et al. (2023); Murali, Zickgraf, et al. (2023). For activity profiling, feces were collected on day 23 for the first sampling group and days 1, 14 and 22 for the second (**Figure 1**).

**Figure 1.**
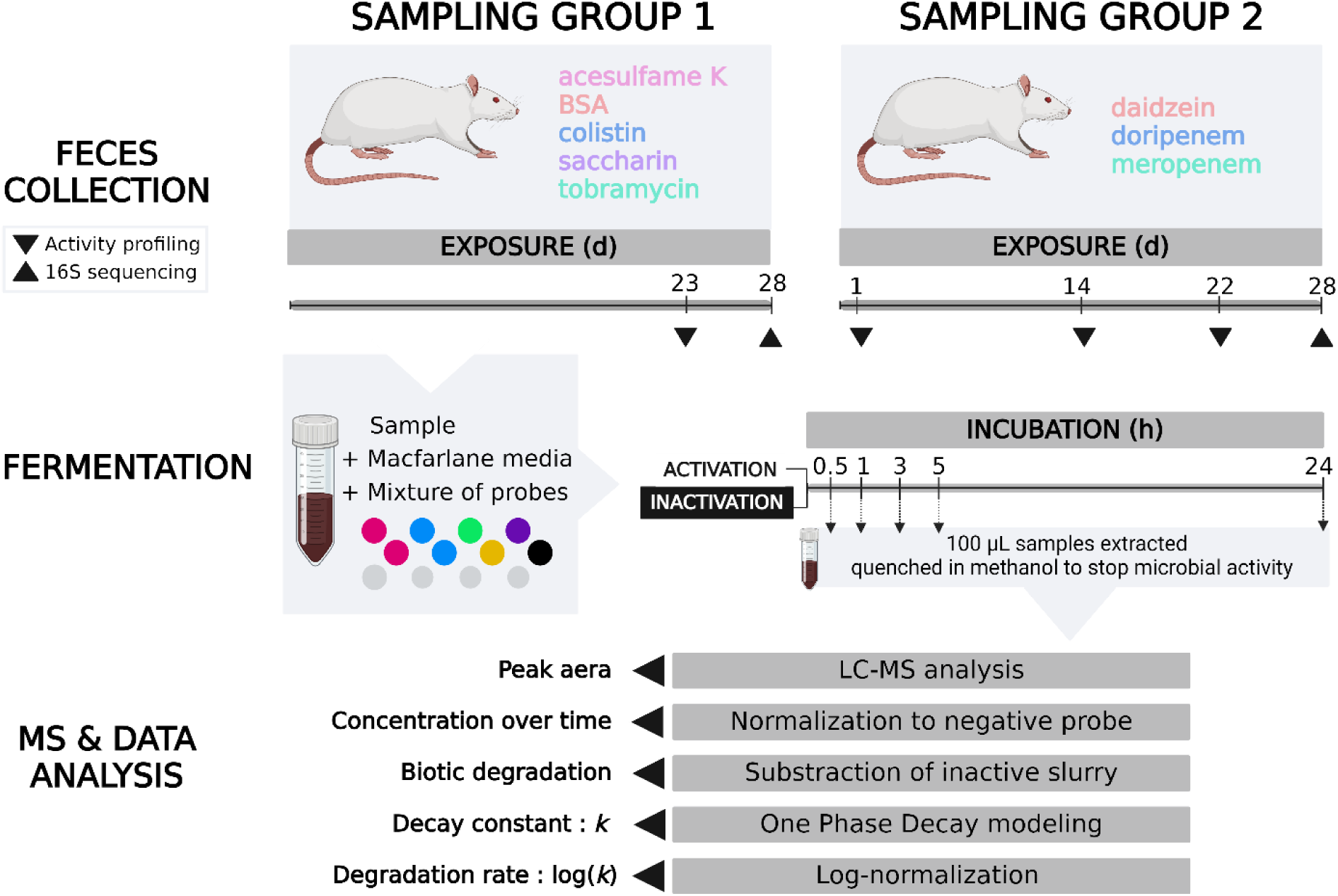
Organization of animal sampling groups, fecal fermentations, and quantification of probe decay rates.

### Fecal slurry preparation

Rat fecal pellets were directly deposited into sterile, pre-weighed microcentrifuge tubes (2 or 5 mL). The headspace was purged with N_2_ and the tubes were kept on ice. Phosphate-buffered saline with 10% (v/v) glycerol was added to produce a 20% (w/v) slurry. The tubes were flushed with N_2_, flash frozen with dry ice, stored at −80 °C until use and shipped on dry ice from the animal experiment facility to the fermentation laboratory. Since the freezing process can induce osmotic and mechanical stresses on bacterial cells, leading to alterations in microbial composition and biotransformation capacity (Aguirre et al., 2015), glycerol was used as a cryoprotectant (Waite et al., 2013). Due to unavoidable technical constraints, the fecal pellets from sampling group 2 were initially stored at −80 °C as fecal pellets, and later suspended in the same manner as sampling group 1. As this deviation in sampling method could potentially influence biotransformation capacity of the microbiota, data analysis for the biotransformation capacity of the two sampling groups was performed separately.

Macfarlane media (Macfarlane et al., 1998) supplemented with volatile fatty acids (Bircher et al., 2018), was prepared under anaerobic conditions in Hungate tubes (Millian), purged with N_2_ gas and autoclaved at 121 °C to ensure sterility (Systec VX-150 autoclave, Systec, Cham, Switzerland). Macfarlane media was chosen to support the growth of maximum microbial taxa and to mimic the richness of gut chyme (Isenring et al., 2023; Macfarlane et al., 1998). Fecal slurries were inoculated at 5% (v/v) into the media within an AtmosBag glove bag (Sigma-Aldrich). This inoculation ratio was chosen due to practical constraints, including the limited availability of rat fecal slurry and the need to maintain the required working volume in Hungate tubes. An anaerobic environment was generated by N_2_ purge, and by use of AnaeroGen anaerobic atmosphere-generating bags (Thermo Scientific) (van Vliet et al., 2019). The absence of oxygen was confirmed by anaerobic indicator test strips (Millipore) and the interior of the bag was sterilized with 80% (v/v) ethanol between the slurry transfers. Following inoculation, the tubes containing the suspended fecal slurries were sealed, removed from the anaerobic environment, and incubated for activation (1 h, 37 °C in a HeraTherm compact microbiological incubator) (**Figure 1**).

### Incubation of probes

A standard mixture of 12 probes (**Figure 2**), in dimethyl sulfoxide (DMSO) and sterile-filtered through a 0.2 µm PVDF filter (polyvinylidene difluoride, BGB analytics) was added to the fecal fermentation mixtures to achieve a final concentration of 1 µM of each probe and a final DMSO concentration of 2% in the mixture. The 12 probes were selected aimed to profile diverse relevant microbial xenobiotic metabolic processes (see **Figure 2**). (Maier et al., 2018; Spanogiannopoulos et al., 2016; Wilson & Nicholson, 2017; Zimmermann et al., 2019), namely nitro reduction, sulfoxide reduction, azo bond reduction, benzisoxazole reduction, amide bond hydrolysis and deglucuronidation (**Figure 2**). Among these compounds, 4 negative probes that are not identified as substrates for gut microbiota metabolism were added for normalization (praziquantel), or comparison (antazoline, disopyramide, tacrine).

**Figure 2.**
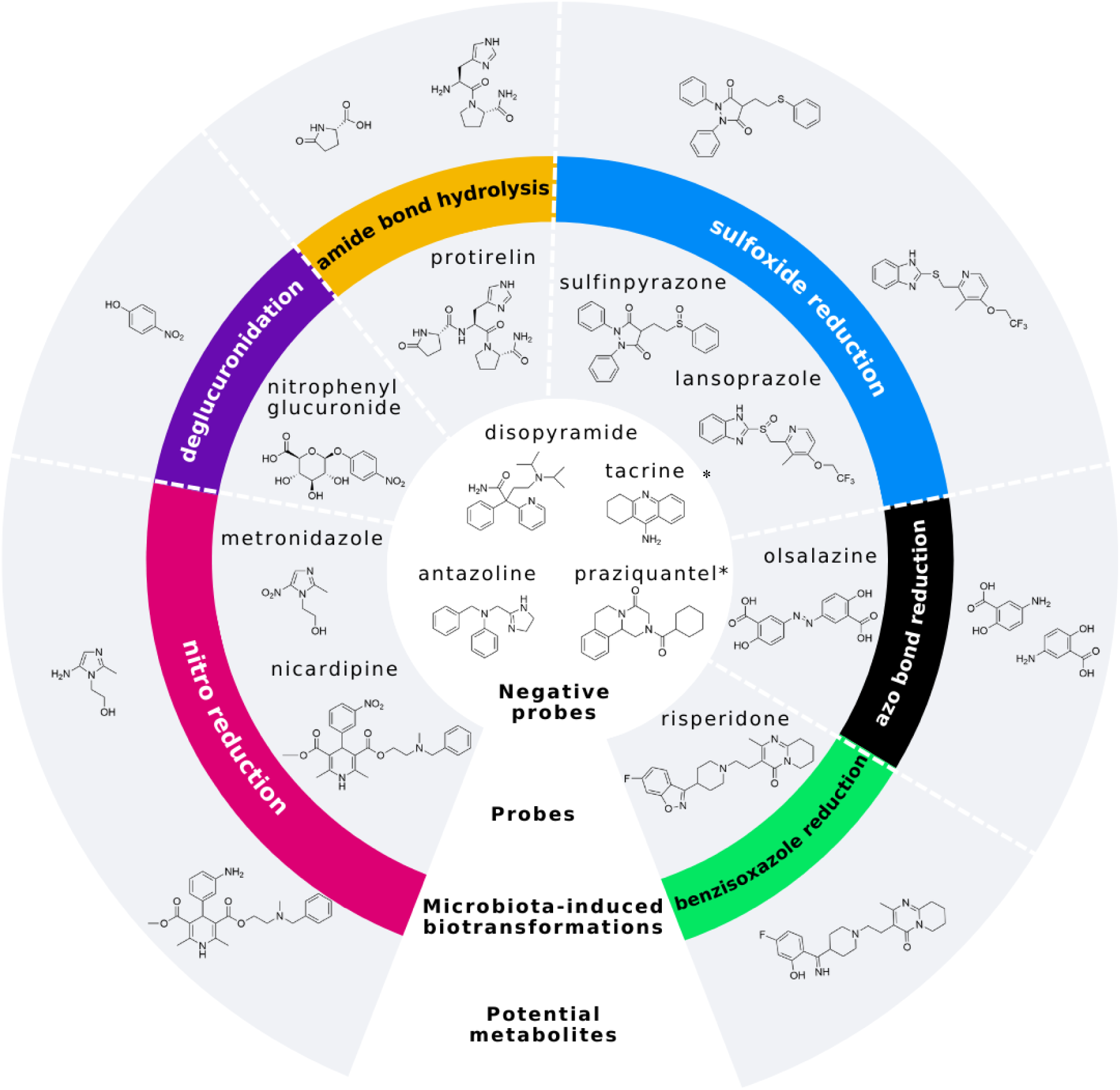
Overview of probes and microbiota-induced reactions profiled herein. *Probe used for normalization.

Following the addition of the probes, the resulting fecal slurry mixtures from exposed and control animals were allowed to incubate in the absence of oxygen for 24 h at 37 °C. To reduce batch effects in data analysis, biological replicates were not incubated in the same batch. Within each batch, the identity of each sample was anonymized until the analysis was completed. Samples of 100 µL were removed from the fermentation mixture at specified times (0.5, 1, 3, 5 and 24 h). The time refers to time elapsed following the addition of probes (**Figure 1**). Maximum incubation time of 24 h avoided factors limiting the accuracy of long-term batch experiments, such as depletion of nutrients, decrease in pH and the accumulation of fermentation- and pH-dependent growth-inhibiting metabolites over time (Isenring et al., 2023). To halt microbial activity and preserve the metabolic state of each sample, they were quenched by the addition of 900 µL of ice-cold LCMS-grade methanol. Additionally, for each incubation, the initial (0 h) sample was obtained by quenching the incubation with ice-cold methanol prior to the addition of probes. The probes were then added, and the sample was processed in the same manner. Quenched samples were stored at −80 °C until analysis by HPLC-HRMS (high-performance liquid chromatography-high resolution mass spectrometry). Furthermore, to determine the abiotic baseline degradation rate of the probes, autoclave-inactivated fecal slurry controls were conducted and processed in the same manner.

### Quantification of probe concentrations by HPLC-HRMS analysis

To determine the rate of degradation of the probes caused by gut microbiome, the change in their concentrations over time was quantified by HPLC-HRMS. Samples were vortex-mixed and centrifugated (4 °C, 20 min, 20,000 x g) using a microtube rotor (Eppendorf). Then, 200 µL of the resulting supernatant was diluted with water (800 µL) containing 50 and 500 nM of analytical internal standards (sulfamethoxazole and 5-fluorouracil, respectively). Prior to analysis, the samples were filtered through a 0.2 µm water-wettable/hydrophilic PTFE (polytetrafluoroethylene) filter. Samples were kept on ice to prevent degradation. HPLC-HRMS analysis was performed on a Thermo Vanquish core HPLC system in line with a Thermo Tribrid ID-X mass spectrometer, operated and monitored using Xcalibur software (version 4.4.16.14). Chromatographic separation was achieved using a Phenomenex Synergi column (4 µm Polar-RP 80 Å, 30 × 2 mm) with a corresponding guard column, changed every 6,000 injections. The HPLC method used solvents A (LCMS-grade water with 0.1% formic acid) and B (LCMS-grade acetonitrile with 0.1% formic acid); gradient 0 min (5% B), 3 min (100% B), 4 min (100% B), 4.1 min (5% B), and 6 min (5% B); flow rate of 0.4 mL/min. The column temperature was maintained at 40 °C, with an autosampler temperature of 4 °C. The injection volume was 10 µL. Heated electrospray ionization was used, with positive (3500 V) and negative (−2500 V) ionization conducted in separate runs. Key parameters were as follows: sheath gas, 50 Arb; auxiliary gas, 10 Arb; sweep gas, 1 Arb, ion transfer tube temperature, 300 °C; vaporizer temperature, 350 °C; expected LC peak width, 3 s. Full scan MS1 was performed using the Orbitrap at 120,000 resolution with a scan range of 120-900 m/z and subsequent data-dependent MS2 acquisition. For MS2, two targeted mass filters were applied with isolation mode quadrupole, 0.4 m/z isolation window, and higher-energy collisional dissociation (HCD) stepped at 30, 45, and 70. Data processing was carried out using QuanBrowser software, involving quantification from MS1 data (10 ppm resolution).

### Data analysis

#### Calculation of probe degradation rates

Probe degradation rates were determined first by normalizing concentrations to the value for the negative probe praziquantel. For all other probes, their level values were expressed as percentage, relative to initial concentration. To account for abiotic (baseline) degradation rates, values for probe levels in autoclave-inactivated fecal slurry controls were subtracted from those derived from active slurries. If not stated otherwise, computations were performed using R (version 4.2.0). Probe degradation rates (*K*) were computed with GraphPad Prism (version 9.2.0, Boston, MA, USA) using the one phase nonlinear decay model. The model, constrained with Y0 = 100 and plateau = 0, employs a one-phase decay (OPD) function to describe processes where a value decreases over time. The OPD function is defined by the equation (1):

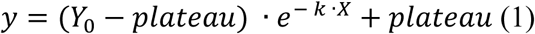

Here, *Y*_*o*_ is the initial value, plateau is the asymptotic value as time (*X*) becomes large, and (*K*) is the decay constant. Degradation rate value is the logarithm base 10 of constant decay. The definition of log(*k*) limits involved setting the log(*k*) values within constraints where 1 ≤ *y* < 100 for *X* = 0.5 h, ensuring that all log(*k*) values fell within this specified interval. Values outside this range were adjusted to these defined limits. The logarithmic fold change (LogFC) was used to compare decay rate between samples groups, according to the equation (2):

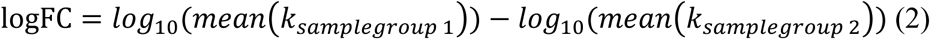

Stringency of statistical testing was verified by false discovery rate (FDR) testing and by application on sufficiently large cohorts (**Table S3**).

#### 16S data analysis

DNA isolation, amplicon-based 16S rRNA gene sequencing and data processing from fecal samples collected on day 28 of both studies were described in (Murali et al., 2022). DNA was amplified using 16S V3-V4 primers (Bakt_341F: 5′-CCTACGGGNGGCWGCAG-3′ and Bakt_805r: 5′-GACTACHVGGGTATCTAATCC-3′). Sequencing was performed on the Illumina MiSeq® next-generation sequencing system (Illumina Inc., San Diego, CA, USA). Signals were processed to fastq files, and the resulting 2 × 250 bp reads were demultiplexed using the MiSeq®-inherited MiSeq Control Software (MCS) v2.5.0.5. As a part of data cleanup, amplicon sequence variants (ASVs) or reads that did not have taxonomic assignment up to the family level were removed. Furthermore, ASVs with non-zero counts in at least two samples were retained, and the others were removed. A quantitative summary of this filtering is available as **Table S4**. Count normalization of the data set was done with cumulative sum scaling, implemented in the publicly available metagenomeSeq Bioconductor package (Paulson et al., 2013). The filtered and normalized data were used for Spearman analysis of correlation with log-normalized *K* values of the probes, per sample. Benjamini and Hochberg multiple hypothesis test correction was applied.

## Results

### Activity profiling with probes indicates alterations of microbial biotransformation capacity

To evaluate the microbial biotransformation capacity of rat fecal samples, we monitored the degradation rates of inoculated probes over 24 h fermentations. The range in degradation rates (values of log(*k*)) of probes spanned from 0.96 (maximal rate possible to measure) to –5.4, reflecting six orders of magnitude range in rates (**Figure 3**). The probe risperidone was degraded the fastest in most samples with relative log(*k*) values ranging between −0.4 and 0.96. We included three probes expected to resist microbial degradation (antazoline, disopyramide, and tacrine) as negative probes, and indeed, their relative degradation was minimal, corresponding to log(*k*) values averaging around –3. Variability in probe degradation rates and the influence of exposure was observed, and three case examples are visualized, depicting probes with fast, slow, or null degradation rates (nicardipine, sulfinpyrazone, and antazoline, respectively) (**Figure 3, A-F**).

**Figure 3.**
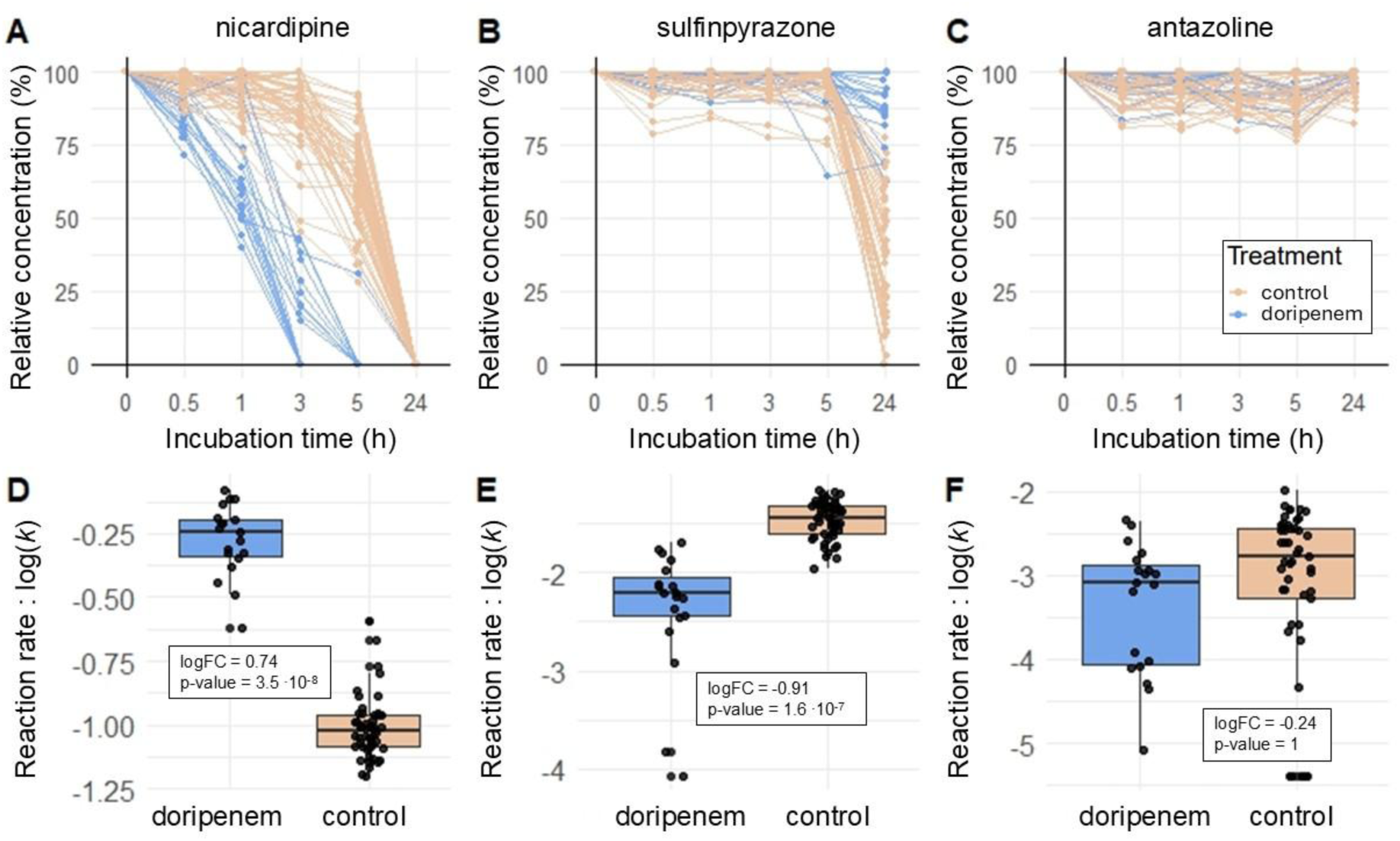
Degradation of three representative probes (A: nicardipine; B: sulfinpyrazone; C: the negative probe antazoline) in ex vivo fermentations of samples from rats exposed with doripenem LD (blue) and non-exposed control (red) animals. Each line represents one fecal sample, and each point represents one sampling time point during ex vivo fermentations.

One particularly interesting example was observed for exposure to doripenem LD, whereby the degradation rate of nicardipine increased and sulfinpyrazone decreased (**Figure 3, A,B,D,E**). More specifically, nicardipine was completely degraded after 3 h in half of the samples from the exposed rats, and after 24 h for all samples, corresponding to log(*k*) values of between 0 and −1.25 (**Figure 3, A, D**), and was significantly increased compared to control rats (p-value = 3.5E-8, Wilcoxon’s rank-sum test, n≥19, **Table S3**). The logFC of 0.74 indicated that the decay was 5.5 times faster in microbiota of exposed rats (**Figure 3 A, D**). In contrast, degradation of sulfinpyrazone was slower, mostly incomplete after 24 h, (log(*k*) of −4.1 to −1.2, **Figure 3, B, E**), and significantly decreased in microbiota of exposed rats (p-value = 1.6E-7, Wilcoxon’s rank-sum test, n≥19, **Table S3**), with a −0.91 logFC, meaning that the degradation of the sulfinpyrazone was 8.1 times slower in microbiota of exposed rats (**Figure 3 B, E**). The probes with the most significant differences in log(*k*) between exposure and control groups were nicardipine and risperidone, with degradation rates altered by five exposure groups out of 16 (**Table 1**). No exposure induced significant differences in degradation rates of the negative probes used in the study (antazoline, disopyramide, and tacrine, **Figure 3 C, F, Table 1**, **Table 2**).

**Table 1.** Logarithmic fold change (LogFC) of mean log(*k*) values from exposed compared to respective controls. Low dose (LD) and high dose (HD) exposure groups are indicated. LogFC of statistically significant differences (Wilcoxon’s rank-sum test, p-value <0.05, n≥8 see Table S3 for details) are indicated in bold text, with background color indicating their sign: green for values < 0 and pink for values > 0. Non-significant differences are indicated by the sign of the logFC.

|  | acesulfame K |  | BSA |  | colistin |  | daidzein |  | doripenem |  | meropenem |  | saccharin |  | tobramycin |  |
| --- | --- | --- | --- | --- | --- | --- | --- | --- | --- | --- | --- | --- | --- | --- | --- | --- |
|  | LD | HD | LD | HD | LD | HD | LD | HD | LD | HD | LD | HD | LD | HD | LD | HD |
| metronidazole | <0 | <0 | >0 | <0 | <0 | >0 | <0 | <0 | **1.02** | **0.76** | **0.91** | **1.14** | >0 | <0 | **1.12** | >0 |
| nicardipine | <0 | <0 | <0 | <0 | >0 | >0 | >0 | >0 | **0.74** | **0.48** | **0.66** | **0.83** | <0 | >0 | **0.50** | >0 |
| sulfinpyrazole | >0 | <0 | >0 | >0 | <0 | <0 | >0 | >0 | - | - | - | - | >0 | >0 | <0 | <0 |
| lansoprazole | <0 | <0 | <0 | >0 | >0 | >0 | >0 | >0 | **0.08** | **0.05** | **0.07** | **0.12** | >0 | <0 | <0 | <0 |
| olsalazine | <0 | <0 | <0 | <0 | >0 | >0 | >0 | >0 | **0.54** | **0.36** | **0.54** | **0.66** | <0 | >0 | >0 | <0 |
| protirelin | <0 | <0 | <0 | <0 | >0 | >0 | <0 | <0 | **0.34** | **0.31** | **0.30** | **0.43** | >0 | <0 | >0 | <0 |
| risperidone | <0 | <0 | <0 | <0 | >0 | >0 | <0 | <0 | >0 | >0 | **0.40** | **0.40** | >0 | <0 | **0.68** | >0 |
| nitrophenyl- | <0 | <0 | >0 | >0 | <0 | <0 | <0 | >0 | <0 | - | <0 | <0 | <0 | <0 | >0 | >0 |
| antazoline | <0 | >0 | <0 | <0 | >0 | >0 | >0 | <0 | <0 | <0 | <0 | <0 | <0 | <0 | >0 | >0 |
| disopyramide | >0 | <0 | >0 | <0 | >0 | >0 | >0 | <0 | <0 | >0 | >0 | <0 | >0 | >0 | >0 | <0 |
| tacrine | <0 | <0 | <0 | >0 | >0 | >0 | >0 | <0 | >0 | >0 | >0 | >0 | >0 | >0 | >0 | >0 |

**Table 2.**
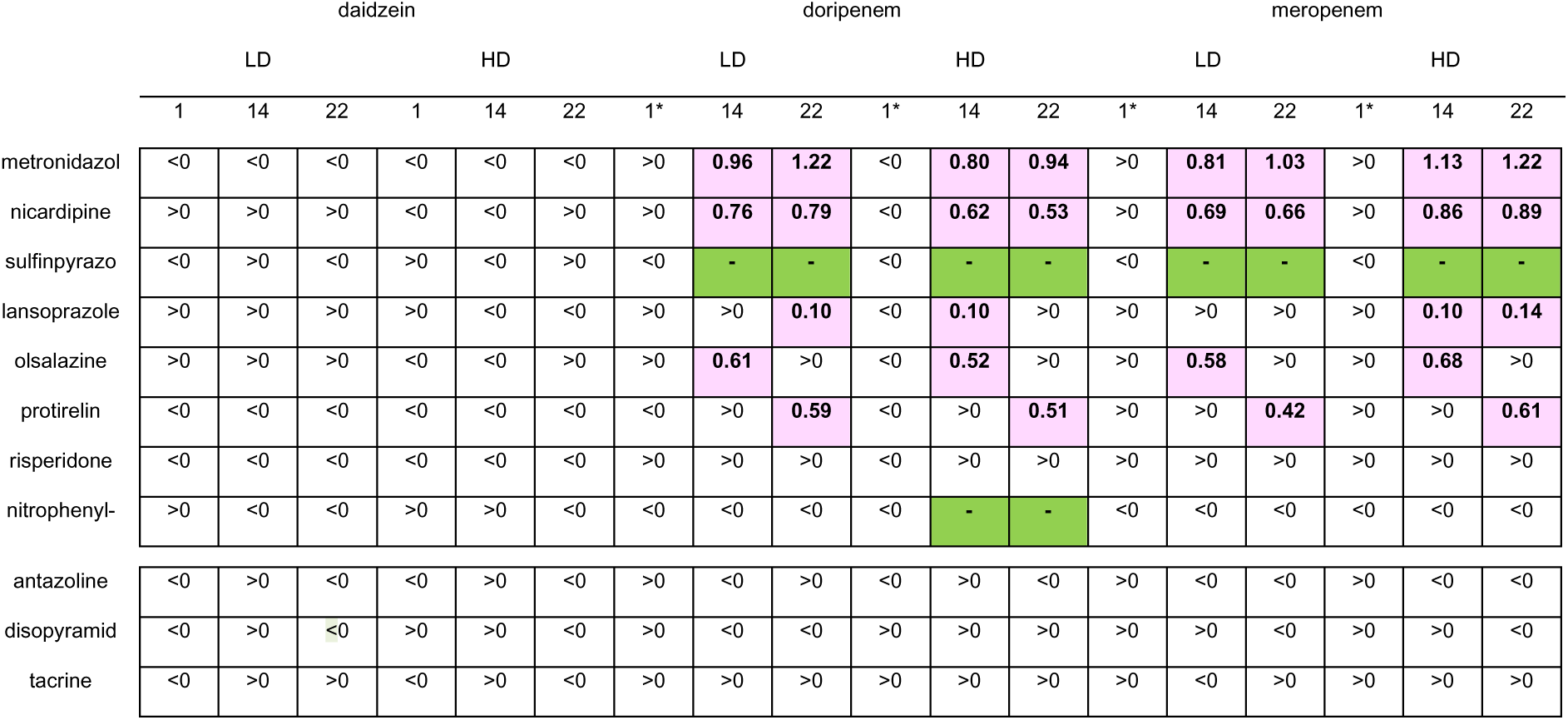
Logarithmic fold change of mean log(*k*) values from exposed compared to respective controls for exposures with multiple timepoints. Low dose (LD) and high dose (HD) exposures groups, as well as sampling days are indicated. LogFC of statistically significant differences (Wilcoxon’s rank-sum test, p-value <0.05, n≥8 see Table S3 for details) are indicated in bold text, with background color indicating their sign: green for values < 0 and pink for values > 0. Non-significant differences are indicated by the sign of the logFC. *statistical analysis were not performed due to limited sample size.

### Alterations of microbiome biotransformation capacities are exposure-specific

We next assessed how treating animals with different chemicals affected the capacity of gut microbiota to degrade probes, comparing exposure groups to each other, and to controls. Based on all probe degradation rates, different exposure groups could be isolated by PCA (**Figure 4**), and clustered on heatmaps of the log(*k*) values (**Figures 5, 6**). Results from sampling groups 1 and 2 were analyzed separately, sex-differences were not considered since no significant correlations were observed (**Tables S5, S6**).

**Figure 4.**
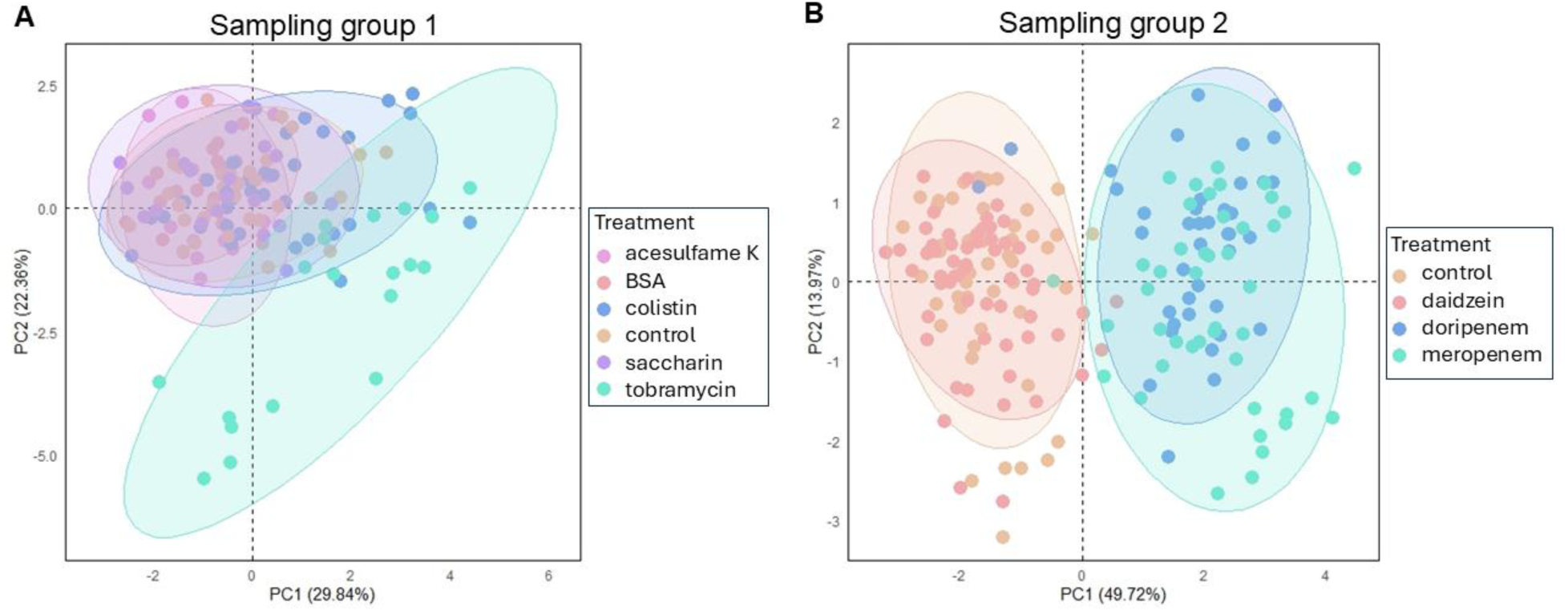
PCA score plot of log(*k*) values for all probes. A. PCA score plot of sampling group 1 samples colored by exposure. B. PCA score plot of sampling group 2 samples colored by exposure. log(*k*) values were normalized to unit variance within each probe. Shaded ellipses represent 95% confidence intervals.

**Figure 5.**
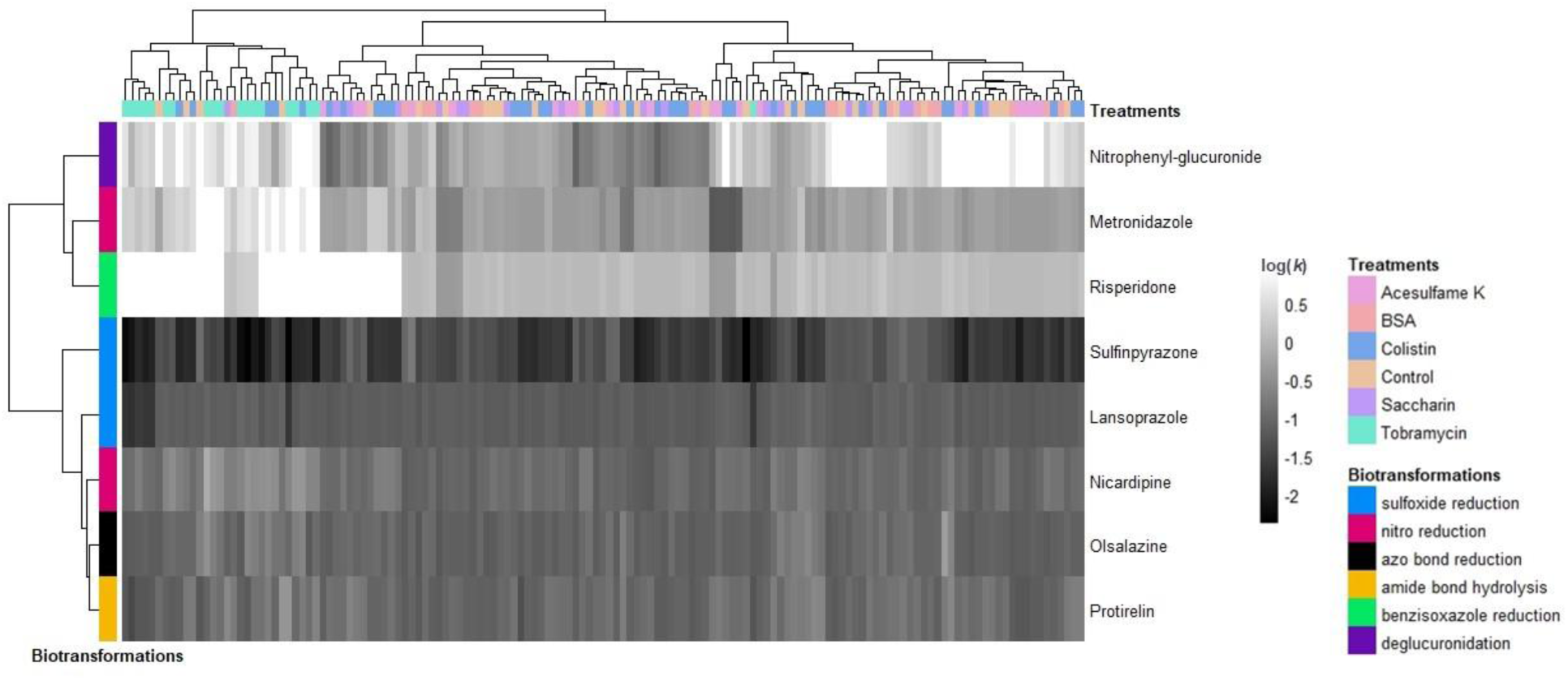
Heatmap of log(*k*) values in sampling group 1, collected on day 23, including hierarchical clustering of positive probes (vertical) and samples (horizontal). Samples were colored according to the exposure of the animal. Probes were colored according to the microbiota-mediated biotransformation type they are expected to undergo.

In sampling group 2, the activity profiles of samples from exposure groups associated with the carbapenem antibiotics meropenem and doripenem were strongly separated from the control and daidzein groups (**Figure 4B, Figure 6**), indicating that these microbiome biotransformation capacities were altered compared to those from the control and daidzein exposure groups. The direction and magnitude of probe degradation rate changes, compared to controls, were similar between doripenem and meropenem, revealing common alterations of microbiome biotransformation capacity induced by carbapenem exposures (**Table 1**). This trend was also consistent across carbapenem dose levels. Specifically, carbapenem exposure decreased the degradation rate of sulfinpyrazone and nitrophenyl-glucuronide (LogFC<0, **Table 1**). In contrast, all carbapenem exposures led to increased degradation rates of metronidazole, nicardipine, lansoprazole, olsalazine, protirelin and risperidone, suggesting an increased functional capacity of the microbiome to transform these probes after exposure-induced microbial alterations. The statistical significance of these exposure-induced alterations in probe degradation rates is detailed in **Table 1**.

**Figure 6.**
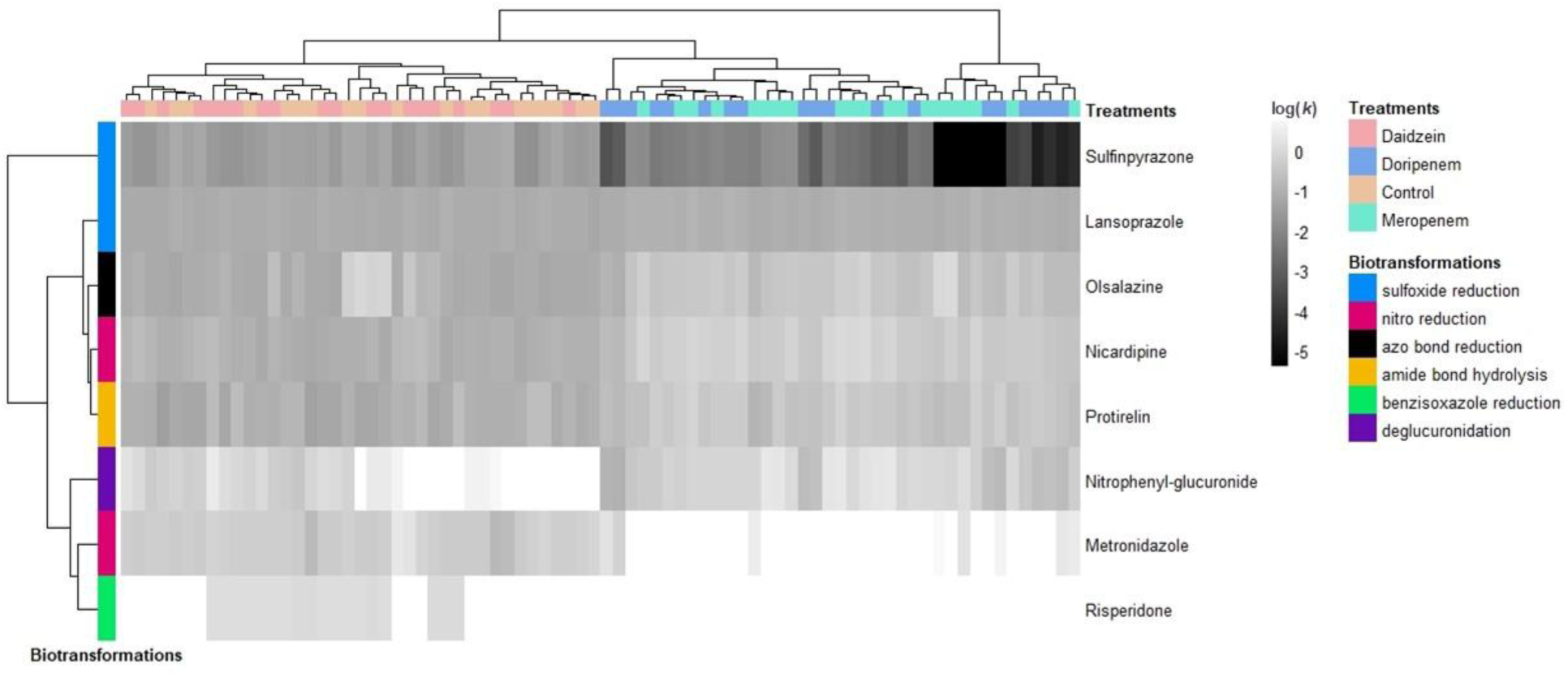
Heatmap of log(*k*) values in sampling group 2 collected on day 22, including hierarchical clustering of positive probes (vertical) and samples (horizontal). Samples were colored according to the exposure of the animal. Probes were colored according to the microbiota-mediated biotransformation type they are expected to undergo.

In sampling group 1, the activity profiles of samples from the tobramycin exposure group strongly separated from the control group (**Figure 4A, Figure 5**), while the colistin exposure group was only partially separated (**Figure 5**), indicating alterations of microbiome biotransformation capacities. Tobramycin and colistin exposure induced alterations of probe activity that were similar to those observed following carbapenem exposures (**Table 1**). In particular, the tobramycin LD exposure significantly increased the degradation rates of metronidazole, nicardipine and risperidone. Exposure groups that did not cluster separately from controls by PCA or hierarchical clustering exhibited no significant differences in LogFC of log(*k*) values (**Figure 4, Figure 5, Table 1**). Thus, microbiome biotransformation capacities were most strongly altered by the antibiotics doripenem, meropenem, and tobramycin LD.

### Biotransformation capacities are altered within the first days of exposure

Further insights into how microbial biotransformation capacities evolved over the 22-day exposure period became evident following exposure to carbapenems (**Figure 1, Table 2, sampling group 2**). Indeed, after only one day of exposure, the degradation rate of some probes was already altered. In general, the direction of alteration in degradation rates was stable across timepoints (except for Doripenem HD), and statistically significant after 14 and 22 days (**Table 2**). The impact of daidzein exposure on the biotransformation capacity of fecal microbiota was nearly negligible for all probes across all sampled days.

### Activity profiling exposes intricate patterns of alterations in microbial biotransformation capacities

Two pairs of probes were associated with the same biotransformation capacity: metronidazole and nicardipine were expected to undergo nitro-reduction, while sulfinpyrazone and lansoprazole were expected to undergo sulfoxide reduction. The degradation of nicardipine and metronidazole followed similar trends, with generally conserved activity profile directions across all exposures compared to control animals (**Table 1, Table 2**). Their degradation rates were significantly increased in microbiomes exposed to doripenem, meropenem, and tobramycin LD. On the other hand, sulfinpyrazone and lansoprazole frequently had logFC’s in opposing directions (**Table 1, Table 2**). The degradation rate of sulfinpyrazone significantly decreased in microbiomes altered by carbapenem exposure, while these same samples exhibited higher capacity to degrade lansoprazole. Additionally, neither of these pairs of probes clustered together by hierarchical clustering of log(*k*) values (**Figure 5, Figure 6**). Interestingly, the arrangement of the probes on the hierarchical clustering dendrograms was nearly identical between the sampling groups (vertical axis, **Figure 5, Figure 6**), but not related to the microbiota-induced biotransformations to which they were expected to be associated (**Figure 2**).

### Alterations in gut microbial biotransformation capacities are correlated with changes in bacterial family abundances

To elucidate relationships between microbiome composition and biotransformation activities, and the impacts of exposure-induced alterations, correlations between abundances of bacterial families, and probe degradation rates were investigated (**Figure 7, Tables S7, S8**). In sampling group 1, the degradation of nicardipine significantly positively correlated with the abundance of Bacteroidaceae (**Figure 7, A**), but no further correlations were evident. For sampling group 2, on the other hand, 105 correlations were observed, and representative relationships are depicted in **Figure 7 (B, C, D, E)**. In all of these correlations, samples from antibiotic-exposed animals could be discriminated from samples from animals exposed with other chemicals. These correlations varied in pattern, with bacterial abundances and degradation rates being higher in either exposure group or similar between them (**Figure 7, B, E, C** or **D**). Finally, the abundances of some bacterial families were very close to the limit of detection in 16S sequencing (e.g., Coriobacteriaceae) and were significantly correlated to probe degradation even with non-detected values for most samples (**Figure 7, E**).

**Figure 7.**
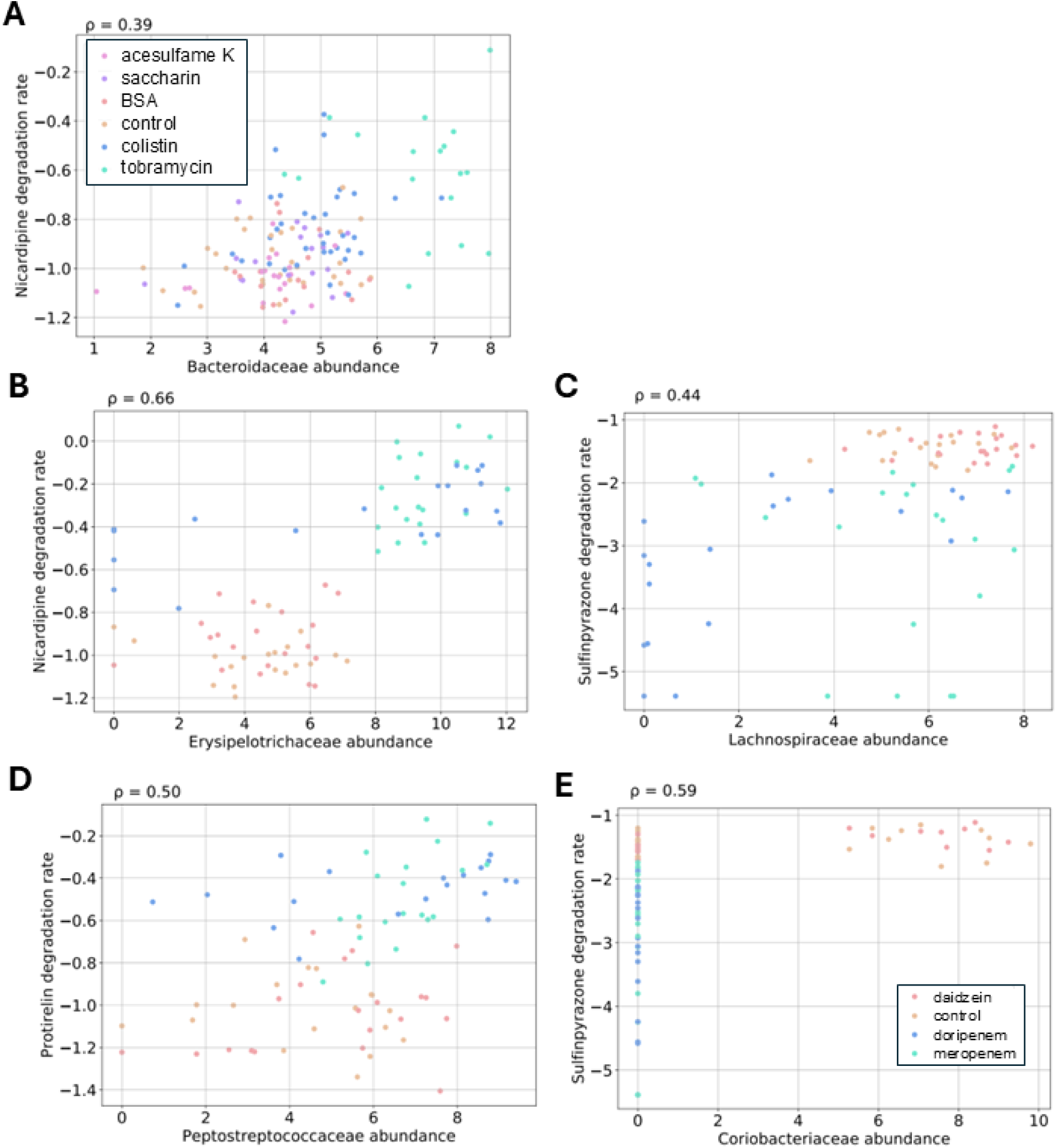
Scatterplots of significant (p < 0.05 FDR) correlations (Spearman analysis) of log(*k*) values versus bacterial family abundances. A. Bacteroidaceae vs nicardipine (sampling group 1). B. Erysipelotrichaceae vs nicardipine (sampling group 2). C. Lachnospiraceae vs sulfinpyrazone (sampling group 2). D. Peptostreptococcaceae vs protirelin (sampling group 2). E. Coriobacteriaceae vs sulfinpyrazone (sampling group 2).

## Discussion

To address microbiota perturbations through alteration of microbiome biotransformation capacity, we herein established a strategy based on anaerobic ex vivo batch fermentations to quantify microbiota-mediated degradations of representative metabolic probes. With this approach, gut microbiome biotransformation capacity of rats that were exposed to various drugs and dietary chemicals was assessed. Fermented fecal samples catalyzed the degradation of eight metabolically active probes, whose reaction rates were influenced by animal exposure to test chemicals. Exposure-specific alterations of microbiome biotransformation capacities were characterized, and alterations were observed within the first days of some chemical exposure (carbapenem antibiotics). Moreover, correlations between changes in bacterial family abundances and alterations of microbiome biotransformation capacity were observed.

We developed a method to assess gut microbial biotransformation capacity and applied it to profile alterations in gut microbiome samples from rats exposed to various drugs and dietary chemicals. Our results confirm the initial hypotheses on which our methodology was based: specifically, we successfully reproduced the degradation of eight probes by the microbiota (Maier et al., 2018; Spanogiannopoulos et al., 2016; Wilson & Nicholson, 2017; Zimmermann et al., 2019), with some of these degradations already established in the human microbiome (sulfinpyrazone, risperidone and metronidazole), (van de Steeg et al., 2018). Conversely, our results also confirm the stability of three negative probes (disopyramide, tacrine and antazoline), which is likewise a key prerequisite for the validity of our approach. While we observed significant exposure-driven alterations of gut-microbial biotransformation capacity (**Table 1, Table 2**), it is difficult to interpret probe-associated activity. Indeed, pairs of probes expected to undergo the same chemical reaction were not similarly degraded (**Figure 5-6**). For instance, sulfinpyrazone and lansoprazole were both expected to undergo sulfoxide-reduction, yet the impact of exposure on the microbiota’s ability to degrade them was in the opposite direction (**Table 1, Table 2**). One explanation may be that different chemical reactions are occurring than predicted, since we measured degradation of starting compound but not formation of products. We did not track metabolite formation, yet we classified biotransformations based on expected metabolism from literature. For example, formation of the metabolite deoxysulfinpyrazone was observed in the ex-vivo fermentation assay of van de Steeg et al. (2018), confirming that the metabolic reaction associated with the probe sulfinpyrazone is the sulfoxide reduction. While lansoprazole’s degradation by the microbiota has been documented by observation of its microbiota-mediated decay over time (Garcia-Santamarina et al., 2024; Maier et al., 2018), empirical evidence of sulfoxide reduction has not been observed. Other possible explanations for these unanticipated differences are enzyme specificity or high influence of individual probe structural features on their biotransformation rates. Specific insights on actual reactions occurring are still needed to confirm the defined biotransformation reactions categories, requiring extensive metabolite identification and tracking outside the present scope. Moreover, the conservation of the arrangement of the probes on the hierarchical clustering dendrograms (**Figure 5-6**) suggests associated biotransformation capacities.

Our results indicate gut microbiota biotransformation capacity was significatively altered by exposure to antibiotics tobramycin, meropenem and doripenem. Biotransformation capacities of the fecal samples from both carbapenem exposure groups were similarly disrupted and suggest increased nitro-reductase capacity (**Table 3**). Additionally, these alterations were partially shared by tobramycin LD. For all other exposures, no difference in biotransformation capacity was observed compared to control samples, and statistical analysis did not reveal an influence of sex on degradation rate logFC (**Tables S5 and S6**). Moreover, no clear impact of sex on microbiota biotransformation capacity could be observed by the clustering of samples of both studies (**Figure S1 and S2**). Additional insights on gut-microbiota functional alterations were provided from the comparison of results from different exposure timepoints. Thus, for most biotransformation reactions, functional alterations induced by carbapenem exposures were evident from the first day of exposure (**Table 2**). Although the sample size for one day of exposure to doripenem and meropenem was insufficient for statistical analysis (n=2), the observed alterations of microbial biotransformation capacity were consistent and significant at later time points (n>8) (**Table 2, Table S3)**.

**Table 3.**
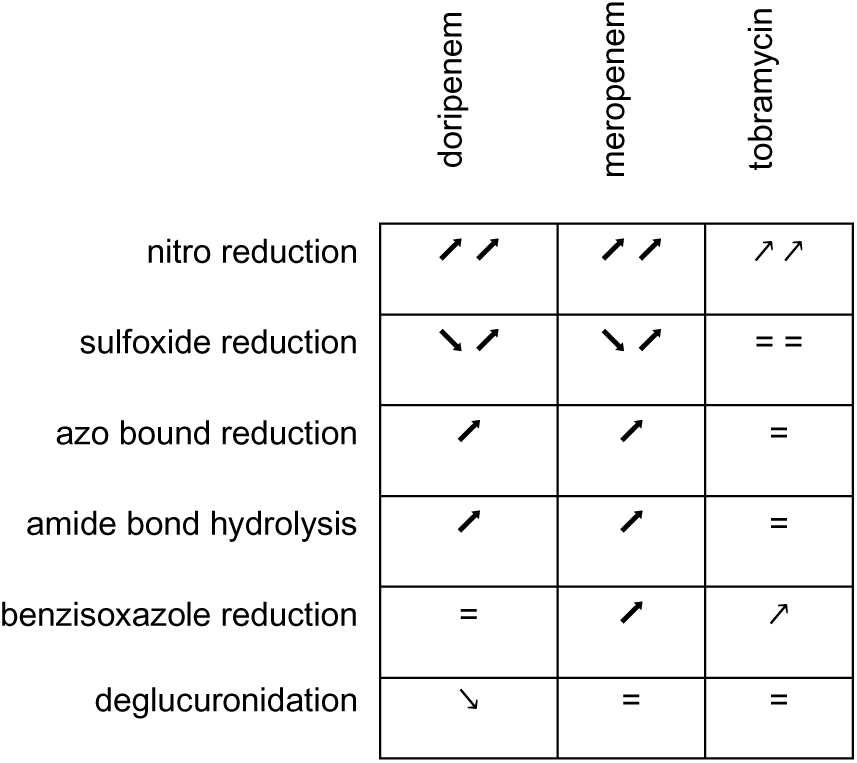
Summary of observed alterations of microbiome biotransformation capacities induced by three antibiotics. Ascending arrows represent an increased degradation rate in exposure compared to control group, descending arrows represent lower degradation rate, equal symbols stand for no significant difference compared to control group. Thick arrows represent significance conserved among both low and high dose groups. The number of symbols per box represents the number of probes associated with a particular biotransformation reaction.

Present findings on the impact of various exposures on gut microbiota biotransformation capacity align with previously observed alterations in intestinal microbiome composition and metabolomes in the same animals (Murali et al., 2022; Murali, Giri, et al., 2023; Murali, Zickgraf, et al., 2023). Significant changes in the gut microbiota and fecal metabolomes of animals exposed to carbapenems were reported, and a significant alteration in bile acid metabolism was particularly highlighted (Murali, Zickgraf, et al., 2023). Additionally, intermediate alteration of microbiota composition was observed from the first day of exposure to carbapenems. Both levels of tobramycin exposure also induced significant alterations in microbial diversity compared to control animals. Metabolite profiling in plasma and feces of tobramycin-exposed animals revealed a strong influence on the metabolome, while colistin had a marginal impact (Murali, Giri, et al., 2023). Artificial sweeteners induced very minor, if any, changes in the gut microbiota and, likewise, limited alterations in fecal metabolite profiles (Murali et al., 2022).

While linking microbiome sequences to biotransformation capacity is a challenge due to the large number of microbial taxa present in intestinal samples, we searched for associations between chemical activities and microbiome signatures, and correlations between bacterial family abundances and probes degradation rates revealed variations in microbial taxa that align with changes in probe transformation rates (**Figure 7**). The low p-values (ranging between 9.6×10^-4^ and 3.9×10^-18^) indicate significant monotonic relationships among our results, representing a plausible link between bacterial family abundance and rate of probe degradation. Sparsity is another important factor that was a driver of significant correlations in this study (**Figure 7E**). The sparsity of values may be due to a lack of sensitivity in sequencing methodology or truly represent lack of presence of certain taxa. Another limitation is that normalizing probe degradation rates to the total sequencing read count per sample introduced outliers due to highly variable read counts (data not shown), and correlations with read-normalized log(*k*) values that primarily reflect bacterial families with abundances near the level of noise, suggesting these results may not be biologically meaningful.

Among the correlated bacterial family-probe pairs (**Table S7 and S8**), chemical activity profiling provided valuable insights into functional disruptions of the gut microbiota, offering a complementary approach to 16S RNA sequencing. While the abundance of bacterial families that were either absent in most samples or unaffected by the exposure did not accurately reflect the microbiota’s state, altered microbiota could still be discriminated by probe degradation rates (**Figure 7 C, D, E**). Thus, activity profiling facilitates effective characterization of altered microbiome biotransformation capacity by reducing the inherent complexity of sequencing-based microbiota composition analysis.

## Conclusion

Chemical activity profiling was used to characterize how oral exposures to drugs and dietary compounds impact rat fecal microbiota biotransformation capacities. Building upon previous reports of alterations in the intestinal microbiota composition and metabolomes of the same animals, we identified corresponding changes in biotransformation capacity associated with in vivo disrupted microbiota communities. Exposure to carbapenem antibiotics impacted the degradation of metabolic probes, increasing the degradation rates of metronidazole, nicardipine, olsalazine and protirelin, while decreasing those of sulfinpyrazone and nitrophenyl-glucuronide. The degradation rates of metronidazole, nicardipine and risperidone were also increased in gut microbiomes from rats exposed to tobramycin. These findings add a layer of understanding to the mechanism by which microbiota disruption may impact host interactions. Furthermore, the activity profiling protocol reported here also enables more functionally relevant study of the impact of exposures on human gut microbiota. This method lays the groundwork for broader applications, and its use in larger, statistically powered cohorts (Ferdous et al., 2022) could extend its relevance to predicting intestinal drug or toxin degradation and exploring the impact of interindividual differences in gut microbiome composition on drug efficacy and chemical toxicity.

## Notes

### Competing Interest Statement

The authors have declared no competing interest.

## References

Aguirre, M., Eck, A., Koenen, M. E., Savelkoul, P. H., Budding, A. E., & Venema, K. (2015). Evaluation of an optimal preparation of human standardized fecal inocula for in vitro fermentation studies. J Microbiol Methods, 117, 78–84. 10.1016/j.mimet.2015.07.019

Appleton, J. (2018). The Gut-Brain Axis: Influence of Microbiota on Mood and Mental Health. Integr Med (Encinitas), 17(4), 28–32.

Bircher, L., Schwab, C., Geirnaert, A., & Lacroix, C. (2018). Cryopreservation of artificial gut microbiota produced with in vitro fermentation technology. Microb Biotechnol, 11(1), 163–175. 10.1111/1751-7915.12844

Bull, M. J., & Plummer, N. T. (2014). Part 1: The Human Gut Microbiome in Health and Disease. Integr Med (Encinitas), 13(6), 17–22.

Carding, S., Verbeke, K., Vipond, D. T., Corfe, B. M., & Owen, L. J. (2015). Dysbiosis of the gut microbiota in disease. Microb Ecol Health Dis, 26, 26191. 10.3402/mehd.v26.26191

Clarke, G., Sandhu, K. V., Griffin, B. T., Dinan, T. G., Cryan, J. F., & Hyland, N. P. (2019). Gut Reactions: Breaking Down Xenobiotic–Microbiome Interactions. Pharmacological Reviews, 71(2), 198. 10.1124/pr.118.015768

Collins, S. L., & Patterson, A. D. (2020). The gut microbiome: an orchestrator of xenobiotic metabolism. Acta Pharmaceutica Sinica B, 10(1), 19–32. 10.1016/j.apsb.2019.12.001

Culp, E. J., Nelson, N. T., Verdegaal, A. A., & Goodman, A. L. Microbial transformation of dietary xenobiotics shapes gut microbiome composition. Cell. 10.1016/j.cell.2024.08.038

Culp, E. J., Nelson, N. T., Verdegaal, A. A., & Goodman, A. L. (2024). Microbial transformation of dietary xenobiotics shapes gut microbiome composition. Cell, 187(22), 6327–6345.e6320. 10.1016/j.cell.2024.08.038

Fan, Y., & Pedersen, O. (2021). Gut microbiota in human metabolic health and disease. Nature Reviews Microbiology, 19(1), 55–71. 10.1038/s41579-020-0433-9

Ferdous, T., Jiang, L., Dinu, I., Groizeleau, J., Kozyrskyj, A. L., Greenwood, C. M. T., & Arrieta, M.-C. (2022). The rise to power of the microbiome: power and sample size calculation for microbiome studies. Mucosal Immunology, 15(6), 1060–1070. 10.1038/s41385-022-00548-1

Garcia-Santamarina, S., Kuhn, M., Devendran, S., Maier, L., Driessen, M., Mateus, A., Mastrorilli, E., Brochado, A. R., Savitski, M. M., Patil, K. R., Zimmermann, M., Bork, P., & Typas, A. (2024). Emergence of community behaviors in the gut microbiota upon drug treatment. Cell, 187(22), 6346–6357.e6320. 10.1016/j.cell.2024.08.037

Isenring, J., Bircher, L., Geirnaert, A., & Lacroix, C. (2023). In vitro human gut microbiota fermentation models: opportunities, challenges, and pitfalls. Microbiome Res Rep, 2(1), 2. 10.20517/mrr.2022.15

Koppel, N., Maini Rekdal, V., & Balskus, E. P. (2017). Chemical transformation of xenobiotics by the human gut microbiota. Science, 356(6344), eaag2770. 10.1126/science.aag2770

Lacroix, C., de Wouters, T., & Chassard, C. (2015). Integrated multi-scale strategies to investigate nutritional compounds and their effect on the gut microbiota. Current Opinion in Biotechnology, 32, 149–155. 10.1016/j.copbio.2014.12.009

Macfarlane, G. T., Macfarlane, S., & Gibson, G. R. (1998). Validation of a Three-Stage Compound Continuous Culture System for Investigating the Effect of Retention Time on the Ecology and Metabolism of Bacteria in the Human Colon. Microb Ecol, 35(2), 180–187. 10.1007/s002489900072

Maier, L., Pruteanu, M., Kuhn, M., Zeller, G., Telzerow, A., Anderson, E. E., Brochado, A. R., Fernandez, K. C., Dose, H., Mori, H., Patil, K. R., Bork, P., & Typas, A. (2018). Extensive impact of non-antibiotic drugs on human gut bacteria. Nature, 555(7698), 623–628. 10.1038/nature25979

Murali, A., Giri, V., Cameron, H. J., Behr, C., Sperber, S., Kamp, H., Walk, T., & van Ravenzwaay, B. (2021). Elucidating the Relations between Gut Bacterial Composition and the Plasma and Fecal Metabolomes of Antibiotic Treated Wistar Rats. Microbiology Research, 12(1), 82–122.

Murali, A., Giri, V., Cameron, H. J., Sperber, S., Zickgraf, F. M., Haake, V., Driemert, P., Walk, T., Kamp, H., Rietjens, I. M. C. M., & van Ravenzwaay, B. (2022). Investigating the gut microbiome and metabolome following treatment with artificial sweeteners acesulfame potassium and saccharin in young adult Wistar rats. Food and Chemical Toxicology, 165, 113123. 10.1016/j.fct.2022.113123

Murali, A., Giri, V., Zickgraf, F. M., Ternes, P., Cameron, H. J., Sperber, S., Haake, V., Driemert, P., Kamp, H., Funk-Weyer, D., Sturla, S. J., Rietjens, I. M. C. M., & van Ravenzwaay, B. (2023). Connecting Gut Microbial Diversity with Plasma Metabolome and Fecal Bile Acid Changes Induced by the Antibiotics Tobramycin and Colistin Sulfate. Chemical Research in Toxicology, 36(4), 598–616. 10.1021/acs.chemrestox.2c00316

Murali, A., Zickgraf, F. M., Ternes, P., Giri, V., Cameron, H. J., Sperber, S., Haake, V., Driemert, P., Kamp, H., Weyer, D. F., Sturla, S. J., Rietjens, I. M. G. M., & van Ravenzwaay, B. (2023). Gut Microbiota as Well as Metabolomes of Wistar Rats Recover within Two Weeks after Doripenem Antibiotic Treatment. Microorganisms, 11(2).

Nicholson, J. K., Holmes, E., Kinross, J., Burcelin, R., Gibson, G., Jia, W., & Pettersson, S. (2012). Host-gut microbiota metabolic interactions. Science, 336(6086), 1262–1267. 10.1126/science.1223813

Noecker, C., & Turnbaugh, P. J. (2024). Emerging tools and best practices for studying gut microbial community metabolism. Nature Metabolism. 10.1038/s42255-024-01074-z

Pascal Andreu, V., Augustijn, H. E., Chen, L., Zhernakova, A., Fu, J., Fischbach, M. A., Dodd, D., & Medema, M. H. (2023). gutSMASH predicts specialized primary metabolic pathways from the human gut microbiota. Nature Biotechnology, 41(10), 1416–1423. 10.1038/s41587-023-01675-1

Paulson, J. N., Stine, O. C., Bravo, H. C., & Pop, M. (2013). Differential abundance analysis for microbial marker-gene surveys. Nature Methods, 10(12), 1200–1202. 10.1038/nmeth.2658

Rooks, M. G., & Garrett, W. S. (2016). Gut microbiota, metabolites and host immunity. Nat Rev Immunol, 16(6), 341–352. 10.1038/nri.2016.42

Spanogiannopoulos, P., Bess, E. N., Carmody, R. N., & Turnbaugh, P. J. (2016). The microbial pharmacists within us: a metagenomic view of xenobiotic metabolism. Nature Reviews Microbiology, 14(5), 273–287. 10.1038/nrmicro.2016.17

Stevanoska, M., Folz, J., Beekmann, K., & Aichinger, G. (2024). Physiologically based kinetic (PBK) modeling as a new approach methodology (NAM) for predicting systemic levels of gut microbial metabolites. Toxicol Lett, 396, 94–102. 10.1016/j.toxlet.2024.04.013

van de Steeg, E., Schuren, F. H. J., Obach, R. S., van Woudenbergh, C., Walker, G. S., Heerikhuisen, M., Nooijen, I. H. G., & Vaes, W. H. J. (2018). An Ex Vivo Fermentation Screening Platform to Study Drug Metabolism by Human Gut Microbiota. Drug Metabolism and Disposition, 46(11), 1596. 10.1124/dmd.118.081026

van Vliet, D. M., Palakawong Na Ayudthaya, S., Diop, S., Villanueva, L., Stams, A. J. M., & Sánchez-Andrea, I. (2019). Anaerobic Degradation of Sulfated Polysaccharides by Two Novel Kiritimatiellales Strains Isolated From Black Sea Sediment [Original Research]. Frontiers in Microbiology, 10. https://www.frontiersin.org/journals/microbiology/articles/10.3389/fmicb.2019.0 0253

Waite, D. W., Deines, P., & Taylor, M. W. (2013). Quantifying the impact of storage procedures for faecal bacteriotherapy in the critically endangered New Zealand Parrot, the Kakapo (Strigops habroptilus). Zoo Biology, 32(6), 620–625. 10.1002/zoo.21098

Wilson, I. D., & Nicholson, J. K. (2017). Gut microbiome interactions with drug metabolism, efficacy, and toxicity. Translational Research, 179, 204–222. 10.1016/j.trsl.2016.08.002

Zierer, J., Jackson, M. A., Kastenmüller, G., Mangino, M., Long, T., Telenti, A., Mohney, R. P., Small, K. S., Bell, J. T., Steves, C. J., Valdes, A. M., Spector, T. D., & Menni, C. (2018). The fecal metabolome as a functional readout of the gut microbiome. Nat Genet, 50(6), 790–795. 10.1038/s41588-018-0135-7

Zimmermann, M., Zimmermann-Kogadeeva, M., Wegmann, R., & Goodman, A. L. (2019). Mapping human microbiome drug metabolism by gut bacteria and their genes. Nature, 570(7762), 462–467. 10.1038/s41586-019-1291-3

