## Supplementary Material for "Chemical activity profiling reveals how exposure to drugs or dietary compounds alters gut microbial biotransformation capacity"

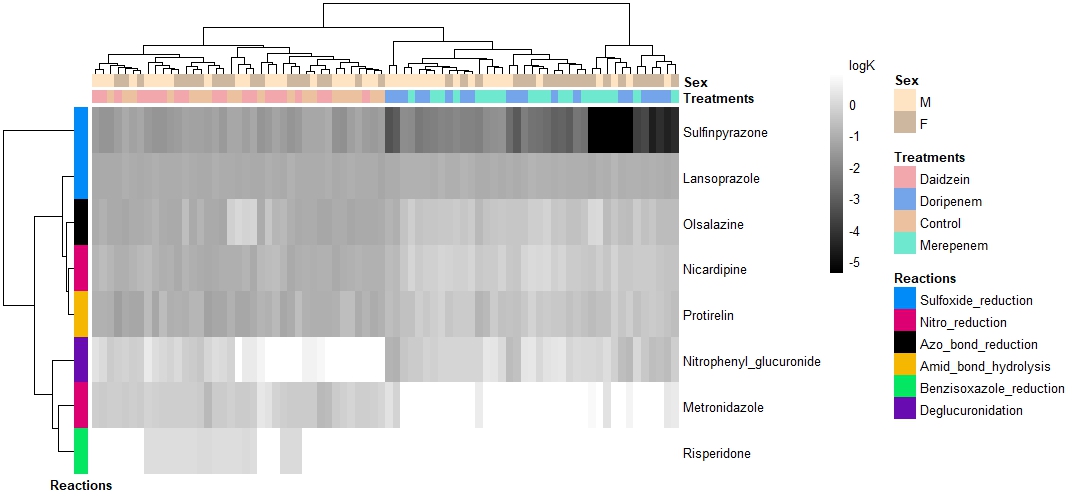
Table S1: Summary and technical information of chemicals used.

| **Chemical** | **CAS** | **Supplier** |
| --- | --- | --- |
| Metronidazole | 443-48-1 | Abcam |
| Risperidone | 106266-06-2 | Fisher scientific |
| 4-nitrophenyl-beta-D-glucuronide | 10344-94-2 | Fisher scientific |
| Simvastatin | 79902-63-9 | Sigma-Aldrich |
| Lovastatin | 75330-75-5 | VWR |
| Antazoline phosphate salt | 154-68-7 | Sigma-Aldrich |
| Tacrine hydrochloride hydrate | 206658-92-6 | Sigma-Aldrich |
| Olsalazine | 6054-98-4 | Sigma-Aldrich |
| Disopyramide phosphate salt | 22059-60-5 | Sigma-Aldrich |
| Terazosin Hydrochloride | 70024-40-7 | Sigma-Aldrich |
| Sulfinpyrazone | 57-96-5 | Sigma-Aldrich |
| Misonidazole | [13551-87-6](https://commonchemistry.cas.org/detail?cas_rn=13551-87-6) | TRC |
| Nicardipine hydrochloride | 54527-84-3 | Sigma-Aldrich |
| Praziquantel | 55268-74-1 | Sigma-Aldrich |
| Sulfamethoxazole | 723-46-6 | Sigma-Aldrich |
| 5-Fluorouracil | 51-21-8 | Fisher scientific |
| Glycerol | 56-81-5 | Sigma-Aldrich |
| Methanol | 69-65-8 | Sigma-Aldrich |

Table S2: Summary of treatments, dose levels and form of preparation of the treatments. All treatments were administered orally by gavage.

| **Sampling group** | **Treatment** | **Low Dose** (mg/kg bw/day) | **High Dose** (mg/kg bw/day) | **Form of preparation** |
| --- | --- | --- | --- | --- |
| 1 | Acesulfame K | 40 | 120 | In deionized water |
|  | BSA | 10 | 100 | In Dulbecco’s phosphate buffered saline |
|  | Colistin | 10 | 100 | In deionized water |
|  | Saccharin | 20 | 100 | In 0,5% carboxymethyl cellulose |
|  | Tobramycin | 100 | 1000 | In deionized water |
| 2 | Daidzein | 100 | 1000 | In corn oil |
|  | Doripenem | 100 | 1000 | In deionized water |
|  | Meropenem | 100 | 300 | In deionized water |

Table S3: Summary of sample stratification.

| **Sampling group** | **Treatment group** | **Dose group** | **Time point** | **n (♀/♂)** |
| --- | --- | --- | --- | --- |
| 1 | Control | - | - | 30 (15/15) |
|  | Acesulfame K | LD | - | 10 (5/5) |
|  |  | HD | - | 10 (5/5) |
|  | BSA | LD | - | 8 (4/4) |
|  |  | HD | - | 8 (4/4) |
|  | Colistin | LD | - | 17 (7/10) |
|  |  | HD | - | 19 (9/10) |
|  | Saccharin | LD | - | 10 (5/5) |
|  |  | HD | - | 10 (5/5) |
|  | Tobramycin | LD | - | 9 (4/5) |
|  |  | HD | - | 10 (5/5) |
| 2 | Control | - | 1 | 10 (5/5) |
|  |  |  | 14 | 20 (10/10) |
|  |  |  | 22 | 19 (9/10) |
|  | Daidzein | LD | 1 | 10 (5/5) |
|  |  | LD | 14 | 10 (5/5) |
|  |  | LD | 22 | 10 (5/5) |
|  |  | HD | 1 | 10 (5/5) |
|  |  | HD | 14 | 9 (5/4) |
|  |  | HD | 22 | 10 (5/5) |
|  | Doripenem | LD | 1 | 2 (1/1) |
|  |  | LD | 14 | 8 (4/4) |
|  |  | LD | 22 | 9 (5/4) |
|  |  | HD | 1 | 2 (1/1) |
|  |  | HD | 14 | 9 (4/5) |
|  |  | HD | 22 | 10 (5/5) |
|  | Meropenem | LD | 1 | 2 (1/1) |
|  |  | LD | 14 | 10 (5/5) |
|  |  | LD | 22 | 10 (5/5) |
|  |  | HD | 1 | 2 (1/1) |
|  |  | HD | 14 | 10 (5/5) |
|  |  | HD | 22 | 10 (5/5) |

Table S4: Summary of 16S data filtering.

| Study | | 1 | 2 |
| --- | --- | --- | --- |
| Number of samples | | 137 | 78 |
| BIOM table | reads | 1’190’944 | 1’617’031 |
|  | ASVs | 20’160 | 2’896 |
| Filtered table | reads | 1’052’251 | 1’391’398 |
|  | ASVs | 1’288 | 904 |

Table S5: Logarithmic fold change (LogFC) of average (mean) log(k) values from treated compared to respective controls for sampling group 1. Low dose (LD), high dose (HD), female (F) and male (M) groups are separated. None of these LogFC was found to be statistically significant (Wilcoxon’s test, p-value <0.05, n in Supplementary table S2). * n<5

| **Treatment** | Acesulfame K | | | | BSA | | | | Colistin | | | | | Saccharin | | | | | Tobramycin | | | |
| --- | --- | --- | --- | --- | --- | --- | --- | --- | --- | --- | --- | --- | --- | --- | --- | --- | --- | --- | --- | --- | --- | --- |
| **Dose** | LD | | HD | | LD | | HD | | LD | | HD | | | LD | | HD | | | LD | | HD | |
| **Sex** | M | F | M | F | M* | F* | M* | F* | M | F | M | F | M | | F | M | F | M | | F* | M | F |
| Metronidazole | -0.25 | -0.29 | -0.23 | -0.35 | -0.05 | 0.25 | -0.10 | -0.02 | 0.13 | -0.15 | 0.02 | 0.06 | 0.26 | | -0.25 | -0.26 | -0.12 | 1.07 | | 1.18 | 0.43 | 0.48 |
| Nicardipine | -0.11 | -0.09 | -0.08 | 0.01 | -0.08 | 0.05 | -0.03 | -0.12 | 0.08 | 0.06 | 0.17 | 0.16 | 0.06 | | -0.11 | 0.00 | 0.02 | 0.46 | | 0.53 | 0.26 | 0.12 |
| Sulfinpyrazone | 0.00 | 0.01 | -0.18 | 0.04 | 0.16 | 0.09 | 0.09 | 0.23 | -0.11 | -0.01 | -0.17 | -0.12 | 0.07 | | 0.06 | -0.09 | 0.21 | -0.34 | | -0.42 | -0.42 | -0.36 |
| Lansoprazole | -0.02 | -0.01 | -0.03 | 0.01 | 0.01 | -0.01 | 0.01 | 0.00 | -0.01 | 0.02 | 0.03 | 0.03 | 0.02 | | -0.02 | -0.01 | -0.04 | -0.02 | | -0.09 | -0.28 | -0.28 |
| Olsalazine | -0.04 | -0.09 | -0.08 | -0.03 | -0.09 | -0.01 | -0.07 | -0.07 | -0.01 | 0.04 | 0.10 | 0.18 | 0.04 | | -0.10 | 0.00 | 0.01 | 0.08 | | 0.31 | -0.01 | -0.08 |
| Protirelin | 0.01 | -0.08 | -0.15 | 0.06 | -0.21 | 0.01 | -0.12 | -0.07 | -0.03 | 0.06 | 0.00 | 0.10 | 0.13 | | -0.05 | -0.04 | 0.01 | 0.07 | | 0.08 | -0.20 | -0.25 |
| Risperidone | -0.07 | -0.24 | 0.03 | -0.04 | -0.28 | 0.05 | -0.31 | 0.01 | 0.22 | 0.10 | -0.03 | 0.19 | 0.23 | | -0.22 | -0.13 | -0.07 | 0.58 | | 0.79 | 0.45 | 0.60 |
| Nitrophenyl_glucuronide | -0.12 | -0.54 | 0.02 | -0.14 | -0.11 | 0.41 | 0.11 | 0.18 | 0.05 | -0.25 | -0.11 | -0.21 | 0.19 | | -0.65 | -0.17 | -0.07 | 0.74 | | 0.22 | 0.50 | -0.07 |
| Antazoline | -0.41 | -0.28 | 0.30 | -0.30 | -0.18 | -0.26 | -0.13 | 0.07 | -0.06 | 0.16 | 0.24 | 0.04 | -0.01 | | -0.43 | -0.73 | -0.04 | 0.14 | | 0.38 | 0.18 | 0.48 |
| Disopyramide | 0.02 | 0.09 | -0.08 | -0.01 | 0.09 | 0.14 | -0.05 | 0.02 | 0.04 | 0.15 | 0.25 | 0.12 | 0.11 | | 0.08 | 0.08 | 0.18 | 0.02 | | 0.25 | 0.02 | -0.06 |
| Tacrine | -1.15 | -0.66 | -0.68 | -0.52 | 0.47 | -0.48 | 0.59 | 0.57 | 0.00 | 0.03 | 0.25 | 0.09 | 0.60 | | 0.02 | 0.18 | 0.43 | 0.48 | | 0.63 | 1.17 | 0.80 |

Table S6: Logarithmic fold change (LogFC) of average (mean) log(k) values from treated compared to respective controls for sampling group 2. Low dose (LD), high dose (HD), sampling days, female (F) and male (M) groups are separated. None of these LogFC was found to be statistically significant (Wilcoxon’s test, p-value <0.05, n in Supplementary table S2). * n<5

| **Treatment** | Daidzein | | | | | | | | Doripenem | | | | | | | | Meropenem | | | | | | | |
| --- | --- | --- | --- | --- | --- | --- | --- | --- | --- | --- | --- | --- | --- | --- | --- | --- | --- | --- | --- | --- | --- | --- | --- | --- |
| **Dose** | LD | | | | HD | | | | LD | | | | HD | | | | LD | | | | HD | | | |
| **Sampling day** | 14 | | 22 | | 14 | | 22 | | 14 | | 22 | | 14 | | 22 | | 14 | | 22 | | 14 | | 22 | |
| **Sex** | M* | F | M | F | M | F | M | F | M* | F* | M | F | M | F* | M | F | M | F | M | F | M | F | M | F |
| Metronidazole | -0.15 | -0.04 | 0.00 | -0.18 | -0.04 | -0.20 | 0.19 | -0.20 | 0.98 | 0.95 | 1.17 | 1.27 | 1.04 | 0.60 | 1.17 | 0.72 | 1.12 | 0.50 | 1.27 | 0.79 | 1.03 | 1.23 | 1.27 | 1.17 |
| Nicardipine | 0.17 | 0.02 | 0.12 | -0.01 | 0.04 | -0.03 | 0.25 | -0.03 | 0.72 | 0.79 | 0.80 | 0.77 | 0.73 | 0.53 | 0.61 | 0.45 | 0.83 | 0.55 | 0.77 | 0.55 | 0.81 | 0.91 | 0.90 | 0.87 |
| Sulfinpyrazone | 0.08 | 0.11 | -0.04 | -0.02 | -0.01 | -0.01 | -0.02 | 0.09 | -0.74 | -0.97 | -1.00 | -0.66 | -2.83 | -2.97 | -2.82 | -2.29 | -0.75 | -0.61 | -0.42 | -2.58 | -2.32 | -0.95 | -1.96 | -1.62 |
| Lansoprazole | 0.13 | 0.00 | -0.01 | 0.02 | -0.02 | 0.01 | 0.02 | -0.02 | 0.08 | 0.05 | 0.07 | 0.12 | 0.12 | 0.08 | 0.05 | 0.04 | 0.07 | 0.07 | 0.06 | 0.04 | 0.09 | 0.10 | 0.16 | 0.12 |
| Olsalazine | 0.16 | 0.01 | 0.21 | -0.09 | -0.05 | -0.01 | 0.17 | 0.04 | 0.59 | 0.63 | 0.52 | 0.57 | 0.60 | 0.45 | 0.41 | 0.26 | 0.67 | 0.49 | 0.59 | 0.44 | 0.62 | 0.73 | 0.74 | 0.64 |
| Protirelin | -0.05 | -0.22 | 0.02 | -0.09 | 0.01 | -0.23 | 0.09 | -0.13 | 0.14 | 0.25 | 0.57 | 0.62 | 0.30 | 0.34 | 0.53 | 0.48 | 0.46 | 0.05 | 0.55 | 0.29 | 0.39 | 0.35 | 0.63 | 0.59 |
| Risperidone | -0.35 | -0.38 | -0.27 | -0.13 | 0.08 | -0.49 | -0.08 | -0.12 | 0.51 | 0.52 | 0.39 | 0.27 | 0.51 | 0.52 | 0.39 | 0.27 | 0.52 | 0.51 | 0.27 | 0.39 | 0.52 | 0.51 | 0.27 | 0.39 |
| Nitrophenyl_glucuronide | -0.01 | -0.43 | 0.03 | -0.04 | 0.18 | -0.04 | 0.10 | -0.12 | -0.35 | -0.45 | -0.24 | -0.71 | -1.03 | -1.11 | -0.92 | -1.28 | -0.06 | -0.34 | -0.36 | -0.41 | -0.72 | -0.45 | -0.60 | -0.17 |
| Antazoline | 0.64 | 0.14 | -0.08 | -0.07 | -0.45 | 0.40 | -0.38 | -0.60 | -0.20 | -0.87 | -0.03 | 0.18 | 0.37 | 0.12 | 0.12 | -0.33 | -0.74 | 0.09 | -0.28 | -0.49 | 0.08 | -0.96 | -0.74 | -1.10 |
| Disopyramide | 0.33 | 0.00 | -0.12 | -0.04 | 0.07 | 0.01 | -0.06 | -0.11 | -0.10 | -0.07 | 0.01 | -0.01 | 0.15 | 0.02 | 0.09 | -0.08 | -0.01 | 0.03 | -0.04 | -0.04 | 0.14 | -0.03 | -0.66 | -0.08 |
| Tacrine | 0.64 | 0.61 | 0.68 | -0.23 | -0.61 | 0.69 | -0.22 | -0.83 | 0.50 | 0.10 | -0.15 | 1.32 | 0.78 | 0.14 | -0.18 | 0.98 | -0.34 | 0.23 | 0.67 | -0.47 | 0.11 | 0.25 | 0.46 | 0.01 |

Table S7: Spearman correlation results between abundances of bacterial families and log(k) values among samples from Sampling group 1. Only correlation results with adjusted p-value (Benjamini & Hochberg false discovery rate) < 0.001 are presented.

| **Probe** | **Bacterial family** | **Correlation** | **p-value** | **FDR adjusted p-value** |
| --- | --- | --- | --- | --- |
| **Nicardipine** |  |  |  |  |
|  | *Verrucomicrobiaceae* | -0.47 | 7.15E-09 | 2.04E-06 |
|  | *Bacteroidaceae* | 0.39 | 2.45E-06 | 2.34E-04 |
| **Metronidazole** |  |  |  |  |
|  | *Peptococcaceae_1* | -0.43 | 1.31E-07 | 1.88E-05 |

Table S8: Spearman correlation results between abundances of bacterial families and log(k) values among samples from Sampling group 2. Only correlation results with adjusted p-value (Benjamini & Hochberg false discovery rate) < 0.001 are presented.

| **Probe** | **Bacterial family** | **Correlation** | **p-value** | **FDR adjusted p-value** |
| --- | --- | --- | --- | --- |
| **Sulfinpyrazone** |  |  |  |  |
|  | *Prevotellaceae* | 0.79 | 3.94E-18 | 3.53E-16 |
|  | *Enterococcaceae* | -0.79 | 4.86E-18 | 3.53E-16 |
|  | *Sutterellaceae* | 0.77 | 2.95E-16 | 8.91E-15 |
|  | *Verrucomicrobiaceae* | 0.76 | 1.24E-15 | 2.81E-14 |
|  | *Desulfovibrionaceae* | 0.74 | 5.58E-15 | 1.07E-13 |
|  | *Rhodospirillaceae* | 0.71 | 2.52E-13 | 3.52E-12 |
|  | *Porphyromonadaceae* | 0.66 | 4.58E-11 | 5.04E-10 |
|  | *Peptococcaceae_1* | 0.66 | 7.30E-11 | 7.79E-10 |
|  | *Anaeroplasmataceae* | -0.63 | 7.06E-10 | 6.75E-09 |
|  | *Rikenellaceae* | 0.63 | 7.73E-10 | 7.20E-09 |
|  | *Coriobacteriaceae* | 0.59 | 1.65E-08 | 1.15E-07 |
|  | *Lactobacillaceae* | 0.53 | 7.72E-07 | 4.12E-06 |
|  | *Lachnospiraceae* | 0.44 | 6.66E-05 | 2.57E-04 |
|  | *Bdellovibrionaceae* | 0.43 | 7.14E-05 | 2.70E-04 |
|  | *Streptococcaceae* | 0.40 | 2.76E-04 | 9.55E-04 |
| **Nicardipine** |  |  |  |  |
|  | *Prevotellaceae* | -0.79 | 4.73E-18 | 3.53E-16 |
|  | *Enterococcaceae* | 0.79 | 7.43E-18 | 4.49E-16 |
|  | *Verrucomicrobiaceae* | -0.77 | 9.20E-17 | 3.71E-15 |
|  | *Sutterellaceae* | -0.77 | 1.20E-16 | 4.37E-15 |
|  | *Rhodospirillaceae* | -0.77 | 1.95E-16 | 6.43E-15 |
|  | *Desulfovibrionaceae* | -0.76 | 4.23E-16 | 1.18E-14 |
|  | *Porphyromonadaceae* | -0.73 | 5.01E-14 | 7.91E-13 |
|  | *Peptococcaceae_1* | -0.72 | 6.32E-14 | 9.55E-13 |
|  | *Rikenellaceae* | -0.70 | 8.25E-13 | 1.07E-11 |
|  | *Erysipelotrichaceae* | 0.66 | 3.42E-11 | 4.00E-10 |
|  | *Clostridiaceae_1* | 0.59 | 1.14E-08 | 8.29E-08 |
|  | *Bifidobacteriaceae* | 0.56 | 8.04E-08 | 4.87E-07 |
|  | *Anaeroplasmataceae* | 0.55 | 1.65E-07 | 9.83E-07 |
|  | *Peptostreptococcaceae* | 0.50 | 2.47E-06 | 1.21E-05 |
|  | *Coriobacteriaceae* | -0.48 | 1.09E-05 | 5.07E-05 |
|  | *Bdellovibrionaceae* | -0.45 | 3.42E-05 | 1.39E-04 |
|  | *Ruminococcaceae* | -0.41 | 2.15E-04 | 7.58E-04 |
| **Olsalazine** |  |  |  |  |
|  | *Porphyromonadaceae* | -0.62 | 1.40E-09 | 1.21E-08 |
|  | *Sutterellaceae* | -0.61 | 2.45E-09 | 2.02E-08 |
|  | *Rhodospirillaceae* | -0.61 | 3.78E-09 | 3.05E-08 |
|  | *Prevotellaceae* | -0.60 | 7.27E-09 | 5.61E-08 |
|  | *Verrucomicrobiaceae* | -0.59 | 1.34E-08 | 9.55E-08 |
|  | *Peptococcaceae_1* | -0.58 | 1.90E-08 | 1.28E-07 |
|  | *Enterococcaceae* | 0.58 | 3.35E-08 | 2.17E-07 |
|  | *Desulfovibrionaceae* | -0.57 | 6.62E-08 | 4.09E-07 |
|  | *Rikenellaceae* | -0.53 | 5.62E-07 | 3.19E-06 |
|  | *Erysipelotrichaceae* | 0.53 | 5.81E-07 | 3.19E-06 |
|  | *Clostridiaceae_1* | 0.47 | 1.50E-05 | 6.80E-05 |
| **Lansoprazole** |  |  |  |  |
|  | *Enterococcaceae* | 0.62 | 1.55E-09 | 1.31E-08 |
|  | *Porphyromonadaceae* | -0.59 | 1.06E-08 | 7.86E-08 |
|  | *Prevotellaceae* | -0.59 | 1.75E-08 | 1.20E-07 |
|  | *Rikenellaceae* | -0.57 | 5.57E-08 | 3.55E-07 |
|  | *Verrucomicrobiaceae* | -0.57 | 6.65E-08 | 4.09E-07 |
|  | *Sutterellaceae* | -0.53 | 4.63E-07 | 2.67E-06 |
|  | *Erysipelotrichaceae* | 0.51 | 1.43E-06 | 7.42E-06 |
|  | *Rhodospirillaceae* | -0.51 | 1.49E-06 | 7.60E-06 |
|  | *Desulfovibrionaceae* | -0.51 | 2.00E-06 | 9.94E-06 |
|  | *Clostridiaceae_1* | 0.46 | 1.99E-05 | 8.59E-05 |
|  | *Peptococcaceae_1* | -0.45 | 3.39E-05 | 1.39E-04 |
|  | *Anaeroplasmataceae* | 0.44 | 4.77E-05 | 1.90E-04 |
|  | *Peptostreptococcaceae* | 0.42 | 1.26E-04 | 4.58E-04 |
| **Risperidone** |  |  |  |  |
|  | *Enterococcaceae* | 0.53 | 5.81E-07 | 3.19E-06 |
|  | *Verrucomicrobiaceae* | -0.51 | 1.63E-06 | 8.24E-06 |
|  | *Desulfovibrionaceae* | -0.45 | 3.66E-05 | 1.48E-04 |
|  | *Sutterellaceae* | -0.44 | 6.74E-05 | 2.57E-04 |
|  | *Peptococcaceae_1* | -0.43 | 1.01E-04 | 3.78E-04 |
|  | *Prevotellaceae* | -0.42 | 1.06E-04 | 3.91E-04 |
| **Metronidazole** |  |  |  |  |
|  | *Enterococcaceae* | 0.83 | 1.19E-20 | 3.90E-18 |
|  | *Sutterellaceae* | -0.78 | 1.88E-17 | 9.74E-16 |
|  | *Prevotellaceae* | -0.78 | 4.77E-17 | 2.16E-15 |
|  | *Peptococcaceae_1* | -0.76 | 5.24E-16 | 1.36E-14 |
|  | *Verrucomicrobiaceae* | -0.76 | 1.36E-15 | 2.90E-14 |
|  | *Rhodospirillaceae* | -0.75 | 1.83E-15 | 3.68E-14 |
|  | *Desulfovibrionaceae* | -0.74 | 8.00E-15 | 1.45E-13 |
|  | *Porphyromonadaceae* | -0.74 | 1.53E-14 | 2.65E-13 |
|  | *Rikenellaceae* | -0.70 | 6.52E-13 | 8.76E-12 |
|  | *Erysipelotrichaceae* | 0.64 | 3.43E-10 | 3.46E-09 |
|  | *Anaeroplasmataceae* | 0.63 | 9.43E-10 | 8.35E-09 |
|  | *Clostridiaceae_1* | 0.58 | 1.98E-08 | 1.31E-07 |
|  | *Coriobacteriaceae* | -0.53 | 7.11E-07 | 3.85E-06 |
|  | *Bifidobacteriaceae* | 0.47 | 1.46E-05 | 6.72E-05 |
|  | *Peptostreptococcaceae* | 0.46 | 2.21E-05 | 9.43E-05 |
|  | *Lactobacillaceae* | -0.41 | 2.34E-04 | 8.16E-04 |
| **Protirelin** |  |  |  |  |
|  | *Enterococcaceae* | 0.82 | 2.15E-20 | 3.90E-18 |
|  | *Verrucomicrobiaceae* | -0.76 | 1.16E-15 | 2.81E-14 |
|  | *Prevotellaceae* | -0.73 | 1.87E-14 | 3.08E-13 |
|  | *Sutterellaceae* | -0.71 | 1.95E-13 | 2.83E-12 |
|  | *Porphyromonadaceae* | -0.68 | 9.97E-12 | 1.25E-10 |
|  | *Rikenellaceae* | -0.67 | 1.58E-11 | 1.91E-10 |
|  | *Desulfovibrionaceae* | -0.66 | 3.91E-11 | 4.43E-10 |
|  | *Anaeroplasmataceae* | 0.64 | 2.75E-10 | 2.85E-09 |
|  | *Peptococcaceae_1* | -0.63 | 5.88E-10 | 5.77E-09 |
|  | *Rhodospirillaceae* | -0.63 | 8.63E-10 | 7.84E-09 |
|  | *Erysipelotrichaceae* | 0.60 | 4.72E-09 | 3.73E-08 |
|  | *Clostridiaceae_1* | 0.60 | 7.42E-09 | 5.61E-08 |
|  | *Peptostreptococcaceae* | 0.50 | 3.10E-06 | 1.50E-05 |
|  | *Bifidobacteriaceae* | 0.49 | 6.15E-06 | 2.94E-05 |
|  | *Coriobacteriaceae* | -0.46 | 2.78E-05 | 1.17E-04 |
|  | *Bdellovibrionaceae* | -0.42 | 1.37E-04 | 4.91E-04 |
| **Nitrophenyl-glucuronide** |  |  |  |  |
|  | *Lachnospiraceae* | 0.53 | 4.53E-07 | 2.65E-06 |
|  | *Rhodospirillaceae* | 0.52 | 1.06E-06 | 5.56E-06 |
|  | *Enterococcaceae* | -0.48 | 9.60E-06 | 4.53E-05 |
|  | *Rikenellaceae* | 0.47 | 1.53E-05 | 6.85E-05 |
|  | *Porphyromonadaceae* | 0.47 | 1.60E-05 | 7.07E-05 |
|  | *Lactobacillaceae* | 0.46 | 1.80E-05 | 7.85E-05 |
|  | *Desulfovibrionaceae* | 0.45 | 2.98E-05 | 1.25E-04 |
|  | *Sutterellaceae* | 0.44 | 5.10E-05 | 2.01E-04 |
|  | *Ruminococcaceae* | 0.44 | 6.70E-05 | 2.57E-04 |
|  | *Anaeroplasmataceae* | -0.42 | 1.10E-04 | 4.02E-04 |
|  | *Prevotellaceae* | 0.42 | 1.50E-04 | 5.33E-04 |

Figure S1: Heatmap of log(*k*) values in study 1, including hierarchical clustering of positive probes (vertical) and samples (horizontal). Samples were colored according to the sex of the animal. Probes were colored according to the microbiota-mediated activity they represent.

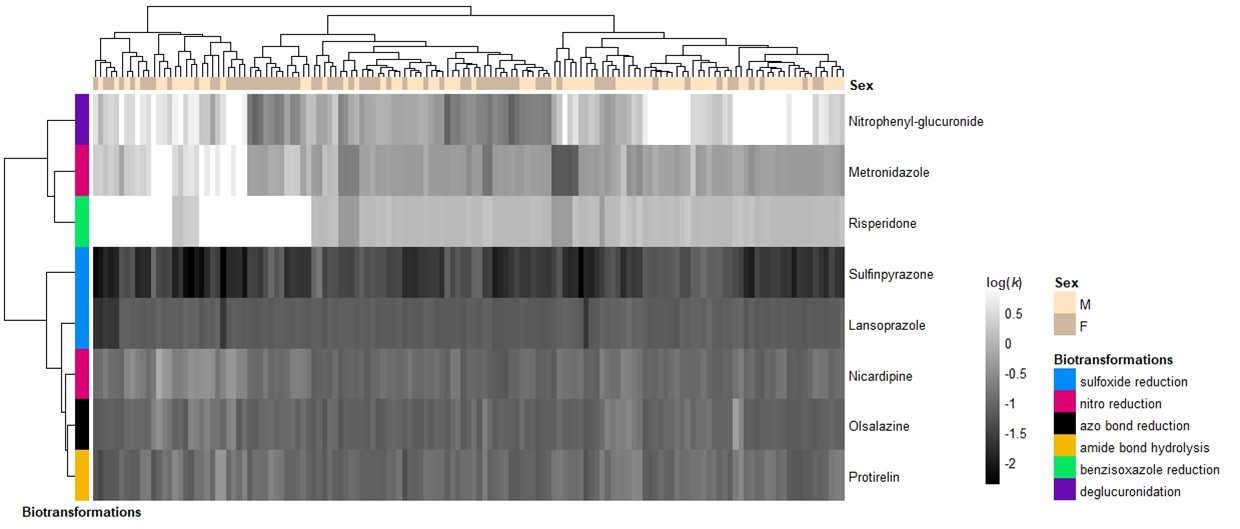

Figure S2: Heatmap of log(k) values in study 2, including hierarchical clustering of positive probes (vertical) and samples (horizontal). Samples were colored according to the sex of the animal. Probes were colored according to the microbiota-mediated activity they represent.

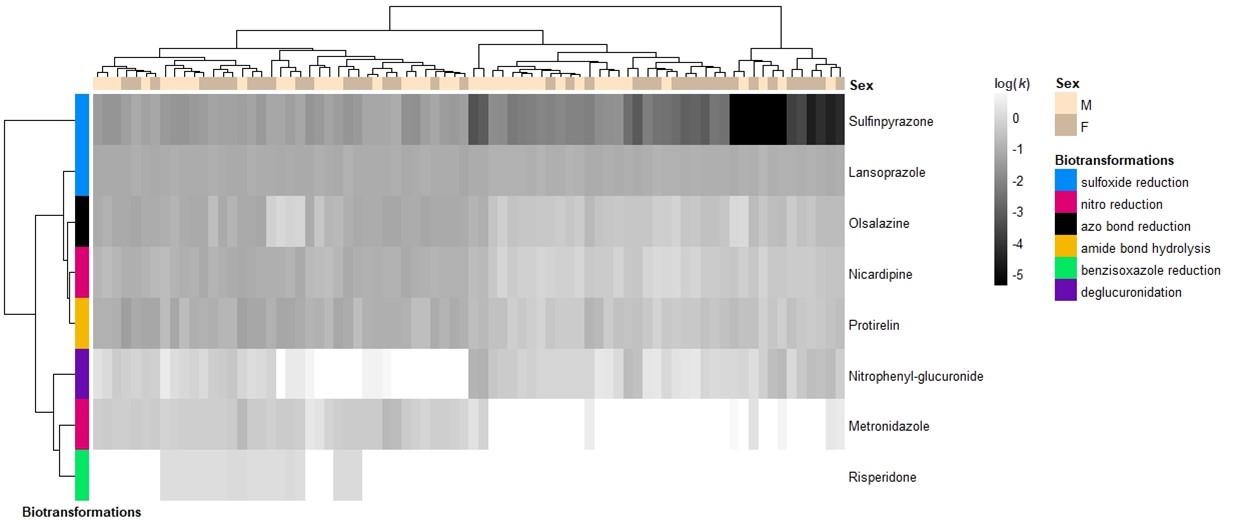
